# Quantifying chromosomal instability from intratumoral karyotype diversity using agent-based modeling and Bayesian inference

**DOI:** 10.1101/2021.04.26.441466

**Authors:** A.R. Lynch, N.L. Arp, A.S. Zhou, B.A. Weaver, M.E. Burkard

## Abstract

Chromosomal instability (CIN) — persistent chromosome gain or loss through abnormal karyokinesis — is a hallmark of cancer that drives aneuploidy. Intrinsic chromosome mis-segregation rates, a measure of CIN, can inform prognosis and are a likely biomarker for response to anti-microtubule agents. However, existing methodologies to measure this rate are labor intensive, indirect, and confounded by karyotype selection reducing observable diversity. We developed a framework to simulate and measure CIN, accounting for karyotype selection, and recapitulated karyotype-level clonality in simulated populations. We leveraged approximate Bayesian computation using phylogenetic topology and diversity to infer mis-segregation rates and karyotype selection from single-cell DNA sequencing data. Experimental validation of this approach revealed extensive chromosome mis-segregation rates caused by the chemotherapy paclitaxel (17.5±0.14/division). Extending this approach to clinical samples revealed the inferred rates fell within direct observations of cancer cell lines. This work provides the necessary framework to quantify CIN in human tumors and develop it as a predictive biomarker.

## INTRODUCTION

Chromosomal instability (CIN) is characterized by persistent whole-chromosome gain and loss through mis-segregation during cell division. Genome instability is a hallmark of cancer (Hanahan and Weinberg, 2011) and CIN is the principal driver of aneuploidy, a feature found in ∼80% of solid tumors (Hancock et al., 2004; Knouse et al., 2017; Weaver and Cleveland, 2006). Importantly, CIN potentiates tumorigenesis (Foijer et al., 2017; Levine et al., 2017; Silk et al., 2013) and associates with therapeutic resistance (Ippolito et al., 2020; Lee et al., 2011; Lukow et al., 2020; Pavelka et al., 2010), metastasis (Bakhoum et al., 2018) and poor survival outcomes (Bakhoum et al., 2011; Denu et al., 2016; Jamal-Hanjani et al., 2017). Thus, CIN is an important characteristic of cancer biology. Despite its importance, CIN has not emerged as a clinical biomarker, in part because it is challenging to quantify.

Although CIN has classically been characterized as a binary variable—tumors either have it or not—recent evidence highlights the importance of the rate of chromosome mis-segregation and the specific aneuploidies it produces. For example, clinical outcomes partially depend on aneuploidy of specific chromosomes (Davoli et al., 2013; Sheltzer et al., 2017; Vasudevan et al., 2020). Further, higher levels of CIN suppress tumor growth when they surpass a critical threshold, thought to be due to lethal loss of essential genes and aberrant gene dosage stoichiometry (Funk et al., 2021; Silk et al., 2013; Weaver and Cleveland, 2008; Zasadil et al., 2014). Moreover, baseline CIN is thought to predict chemotherapeutic response to paclitaxel (Janssen et al., 2009; Swanton et al., 2009) and is predicted to promote detection or evasion from the immune system (Davoli et al., 2017; Santaguida et al., 2017). Clinical determination of a tumor’s intrinsic rate of chromosome mis-segregation will enable validation of CIN as a biomarker for cancer progression and treatment response.

Many approaches have been used to characterize CIN, but struggle to quantify its rate. These include histologic analysis of mitotic defects (Bakhoum et al., 2011; Jin et al., 2020), fluorescence *in-situ* hybridization (FISH) with probes to detect individual chromosomes (Thompson and Compton, 2008), and gene-expression methodologies like CIN scores (Carter et al., 2006). While these methods are readily accessible, they have significant drawbacks for clinical application. FISH and mitotic visualization approaches are laborious. Direct visualization of mitotic defects to measure CIN is only possible in the most proliferative tumors where a sufficient number of cells are captured in short-lived mitosis. For FISH, a subset of chromosomes is quantified, which will be misleading if there is bias toward specific chromosome gains/losses (Dumont et al., 2019). While gene expression scores are proposed as an indirect measure of CIN, they are not specific to CIN and are known to also correlate highly with proliferation signatures and structural aneuploidy (Carter et al., 2006; Sheltzer, 2014).

By contrast, single-cell sequencing promises major advances in quantitative measures of CIN by displaying cell-cell variation for each chromosome across hundreds of cells (Navin et al., 2011; Wang et al., 2014). However, selection poses another complication. Previous single-cell analyses have identified surprisingly low cell-cell karyotype variation, even when mitotic errors are readily observable via microscopy (Bolhaqueiro et al., 2019; Gao et al., 2016; Kim et al., 2018; Nelson et al., 2020; Wang et al., 2014). These observations highlight the confounding role of karyotype selection in measuring CIN in human tumors. Indeed, karyotype selection reduces karyotype variance in cancer cell populations, even after exhibiting mitotic errors (Gerstung et al., 2020; Ippolito et al., 2020; Lukow et al., 2020). It may be possible to overcome this limitation by modeling chromosomal instability and explicitly considering the evolutionary selection of aneuploid cells.

We developed a quantitative framework to measure CIN by sampling population structure and cell-cell karyotypic variance in human tumors, accounting for selection on aneuploid karyotypes. We built our framework on the use of phylogenetic topology measures to quantify underlying evolutionary processes (Mooers and Heard, 1997); in this case to quantify CIN from both the diversity and the aneuploid phylogeny within a tumor. Using an agent-based model of CIN, we determined how distinct types and degrees of selective pressure shape the karyotype distribution and population structure of tumor cells at different rates of chromosome mis-segregation. We then used this *in silico* model as a foundation for parameter inference to provide a quantitative estimate of CIN as a numerical rate of chromosome mis-segregation per cell division. We applied this model to quantify CIN caused by the chemotherapeutic paclitaxel in culture, then, using existing single-cell whole-genome sequencing data (scDNAseq), we quantified CIN in cancer biopsy and organoid samples. As a whole, this work provides the necessary framework to quantify CIN in human tumors as a scalar, and to develop it as a prognostic and predictive biomarker.

## RESULTS

### A framework for modeling CIN and karyotype selection

To assess intratumoral CIN via cell-cell karyotype heterogeneity, we considered how selection on aneuploid karyotypes impacts observed chromosomal heterogeneity within a tumor. By modeling fitness of aneuploid cells, we can observe chromosomal variation in a population of surviving cells. The selective pressure of diverse specific aneuploidies on human cells has not been, to our knowledge, directly measured. Therefore, we employ previously developed models of selection

In transient CIN models, fit karyotypes are selected while unfit karyotypes are eliminated over time (Ippolito et al., 2020; Ravichandran et al., 2018; Sheltzer et al., 2017; Vasudevan et al., 2020). We use two previously proposed models of aneuploidy-associated cellular fitness, as well as a hybrid model, to an agent-based mitosis and mis-segregation framework. The Gene Abundance model is based on the relatively low incidence of aneuploidy in normal tissues and assumes cellular fitness declines as the cell’s karyotype diverges from a balanced euploid karyotype (i.e. 2N, 3N, 4N)(Sheltzer and Amon, 2011; Zhu et al., 2012). When an individual chromosome diverges from euploid balance, its contribution to cellular fitness is weighted by its abundance of genes (Supplemental Figure 1A, left). Alternatively, the Driver Density model assumes that each chromosome contributes to cellular fitness, weighted by its ratio of Oncogenes and Essential genes to Tumor suppressor genes (TOEs)(Davoli et al., 2013; Laughney et al., 2015). For example, selection will favor loss of chromosomes with tumor suppressors and favor gain of chromosomes replete with oncogenes and essential genes (Supplemental Figure 1A, right). The hybrid averaged model accounts for both karyotypic balance and TOE densities (Supplemental Figure 1A, middle). Using these fitness models, we assigned chromosome scores to reflect each chromosome’s value to cellular fitness under each model (Figure 1A), the sum of which represent the total fitness value for the cell, relative to a value of 1 for a euploid cell (Figure 1A). Further, we scaled the impact of selective pressure in these models with an exponent, S, ranging from 0 (no selection) to 200 (high selection). While we recognize these models are approximations, they are nevertheless useful to estimate how mis-segregation and selective pressure cooperate to mold karyotypes in the cell population.

**Figure 1.**
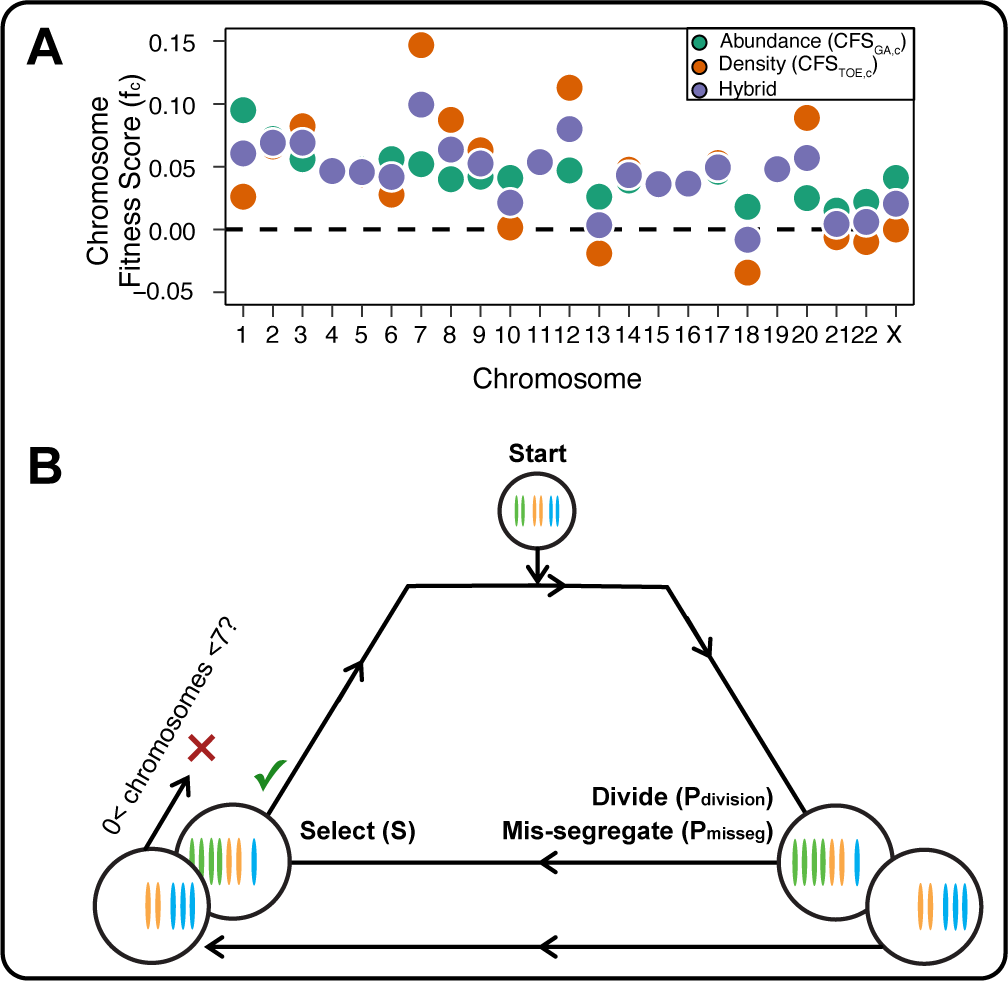
A **framework for modeling CIN and karyotype selection** (A) Chromosome scores for each model of karyotype selection. Gene Abundance scores are derived from the number of genes per chromosome normalized to the number of all genes (Materials & Methods). Driver Density scores come from the pan-cancer chromosome scores derived in Davoli et al. (2013), and normalized to the sum of scores for chromosomes 1-22. The X chromosome did not have a score and was set to 0. Hybrid model scores are set to the average of the Driver and Abundance models. (B) Framework for the simulation of and selection on cellular populations with CIN. Cells probabilistically divide (Pdivision = 0.5) and probabilistically mis-segregate chromosomes (Pmisseg = [0.001… 0.5]). After, cells experience selection under one of the selection models, altering cellular fitness and the probability (Pdivision) a cell will divide again (green check). Additionally, cells wherein the copy number of any chromosome falls to zero or surpasses 6 are removed (red x). After this, the cycle repeats.

Next, we developed an agent-based model of population growth with each cell having its own karyotype. For 40 generations—80 time steps, each being about ½ generation we simulated population growth under a range of selective pressure exponents (S) and rates of chromosome mis-segregation (P_misseg_). Each of the 100 euploid founder cells have a 50% chance of dividing per time step (to partially mimic the asynchrony of cell division; initial P_division_ = 0.5; Figure 1B, Supplemental Figure 1C). When a cell undergoes division, each of the original 46 chromosomes segregate or mis-segregate probabilistically (P_misseg_). After division, the new karyotype of each daughter cell is assessed, and fitness (F) is recalculated according to the fitness model prior to scaling by the selection exponent (S). Further, aneuploidy is lethal if a daughter completely lacks any one chromosome, or exceeds 6 copies of any chromosome, and the cell ‘dies’ and is removed from the population. Any remaining viable cell has its P_division_ adjusted by the cell’s fitness under selection (FS), such that more fit cells are more likely to divide again, contributing more greatly to the cell population. Due to computational limits, cell populations are capped at 4800; when the population surpasses this limit, half are removed at random. In this manner, the model replicates the process by which CIN creates karyotypic heterogeneity in a tumor. Further, it allows random sampling of karyotypes, mimicking single-cell sequencing of a tumor, to study how these dynamics contribute to karyotype evolution and to determine the necessary number of single cells that need to be sampled for a clinical biomarker.

### Evolutionary dynamics imparted by CIN

To understand the interplay between CIN and selection on the cell population, we simulated populations with CIN under each model of karyotype selection for 80 time steps. To reproduce diverse circumstances, we varied the rate of mis-segregation (P_misseg,c_ ∈ {0.001-0.5 per chromosome}; or 0.046 – 23 mis-segregations per cell division) and levels of the selection pressure exponent (S ∈ {0-200}; ranging from no selection to any aneuploidy heavily selected for/against). The probability of division (P_division_) is adjusted by selection for aneuploid cells but starts at 0.5 for the euploid founder population, allowing euploid cells to divide once every 2 time steps on average (Figure 2A). Over time, the simulated cell number rapidly increases to the cap of 4800, whereupon a random half of the population will be deleted. Thus, a population with F^S^ = 1 will remain at about 4800 cells over time (Figure 2B). We measured diversity as the mean of variances for each chromosome across the population and normalized this to the mean ploidy of the population (mean karyotypic variance, MKV). We then plotted heatmaps showing the dynamics of diversity across time, mis-segregation rate, and selection levels (Figure 2C). When mis-segregation rate is increased (y-axis), MKV tends to increase over time (x-axis). Extremely high mis-segregation rates in the absence of selection (S=0) can result in complete population collapse (white area) due to the high incidence of death-triggering copy number states (1>n_c_>6). The three selection models (Abundance, Driver, Hybrid) are displayed as three columns (Figure 2C). As expected, when selection is zero (S=0) the three selection models returned comparable MKV profiles over time. When selective pressure is applied (S>0), the diversity profiles diverge. As expected, the abundance model negatively selects against all aneuploid karyotypes and yields low heterogeneity at all time steps, though it does scale to a low degree with mis-segregation rate. With the Driver model, there is a sharp increase in heterogeneity at moderate mis-segregation rates and the model was more tolerant of high degrees of aneuploidy, to the point where the cell population collapsed due to nullisomy even under moderate selection (S=10). As expected, the Hybrid model fell between that of the Abundance and Driver models because selection only partly favors some aneuploid karyotypes while selecting against most others. Under heavy selection (S=200), the ability for any model to remain significantly diverse is dampened except for a moderate range of mis-segregation rates in the Driver model (Figure 2C).

**Figure 2.**
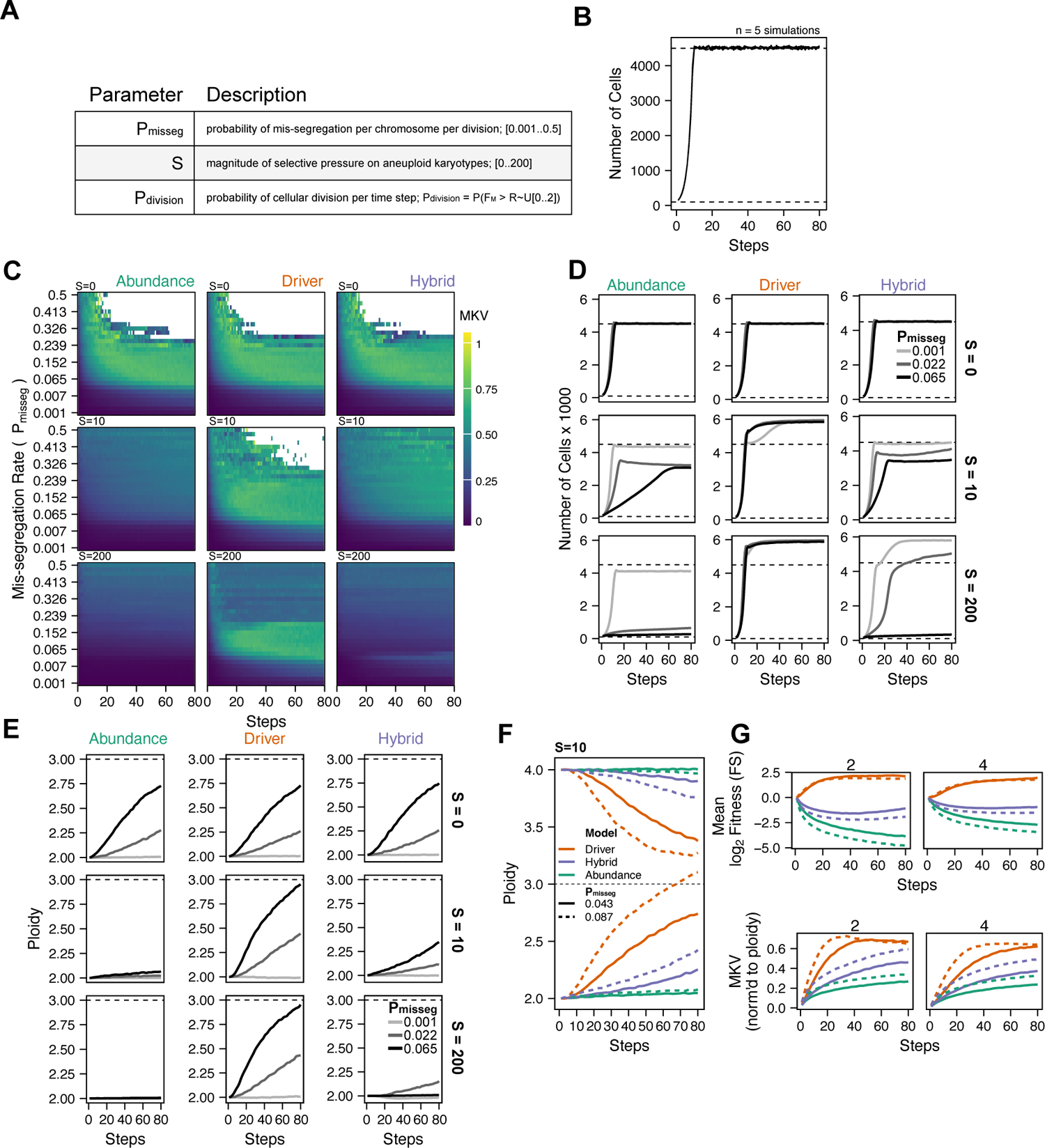
Evolutionary dynamics imparted by CIN. (A) Parameters used to control simulation of populations with CIN. All simulations were performed with an initial Pdivision = 0.5. Thus, 2 steps ∼ 1 generation. (B) Population growth curve in the absence of selective pressure (Pmisseg = 0.001, S = 0, n = 5 simulations). The steady state population in null selection conditions is 4500 cells. (C) Heatmaps depicting dynamics of karyotype diversity as a function of time (steps), mis-segregation rate (Pmisseg), and selection (S) under each model of selection. Columns represent the same model; rows represent the same selection level. Mean karyotype diversity (MKV) is measured as the variance of each chromosome averaged across all chromosomes 1-22, and chromosome X. Low and high MKV are shown in blue and yellow respectively. White space indicates simulations with populations < 5 cells (n = 5 simulations for every combination of parameters). (D) Population growth under each model, varying Pmisseg and S. Pmisseg = [0.001, 0.022, 0.065] translate to about 0.046, 1, and 3 mis-segregations per division respectively. (E) Dynamics of the average ploidy (total # chromosomes / 23) of a population while varying Pmisseg and S. (F) Dynamics of ploidy under each model for diploid and tetraploid founding populations. Pmisseg = [0.043, 0.0.087] translate to about 2 and 4 mis-segregations per division respectively. (G) (Top) Fitness (log2 F^S^) over time for diploid and tetraploid founding populations evolved under each model. (Bottom) Karyotype diversity dynamics for diploid and tetraploid founding populations. MKV is normalized to the mean ploidy of the population at each time step. Plotted lines in D-G are local regressions of n=5 simulations.

We also quantified the population of viable cells (Figure 2D, Supplemental Figures 2B,D). Using the Abundance model at increased levels of selection, the population of cells took longer to grow compared with other models. By contrast, those grown under the Driver model proliferated more rapidly. Under selection, cells grown under the Hybrid model proliferated rapidly at low rates of mis-segregation, while higher rates still limited growth. Additionally, while a random half of any population that reaches 4800 cells is deleted, populations with an average fitness > 1 will proliferate more rapidly than they are deleted, thus these fit populations will exceed the population cap (Figure 2D, Supplemental Figures 2B,D). We noted that in some cases, the mean ploidy of the populations would increase over time (Figure 2E). Without selection (S=0; top), total ploidy increased over time in all models, likely due to chromosome gains being permitted more than losses in our model (since cells ‘die’ with nullisomy or any chromosome > 6, the optimum is 3.0). Once selection is applied (S>0), the models diverge. As proliferation decreases under the abundance model, there is less increase in average ploidy. Alternatively, under the Driver model, the population mean a triploid more rapidly than under selection-null conditions. This is consistent with previous findings using models built on chromosome-specific driver densities (Davoli et al., 2013; Laughney et al., 2015). Under the Hybrid model, with high selection (S=200), the mean ploidy of the population increased only with moderate mis-segregation. This indicates that there is an optimum mis-segregation rate to enable favorable selection of the TOE drivers, balanced against the adverse effects of high mis-segregation rates on the population. Taken together these data demonstrate how selection and mis-segregation rate interact in a complex manner to shape the array of aneuploidy karyotypes found in a population of cells, or a human tumor. Further, they demonstrate that sampling cells and measuring karyotypic diversity in a tumor does not allow direct determination of mis-segregation rates, as diversity is influenced by other factors.

Genome doubling is an event that is thought to occur early, relative to other copy number changes, in the genesis of some tumors (Bielski et al., 2018; Gerstung et al., 2020). Tumors are hypothesized to leverage genome doubling to buffer against loss of chromosomes and thereby favor aneuploidy. To determine how genome doubling impacts evolution in our model, we repeated these models, comparing diploid and tetraploid founders (Figure 2F). Both tend to converge towards the near-triploid state (ploidy ∼ 3.0), as observed in many human cancers (Carter et al., 2012), though this occurs most rapidly with the Driver model. As predicted, average fitness (F^S^) of diploid and tetraploid founding populations differs, with tetraploid cells being buffered against the negative effects of aneuploidy (higher F^S^ values), particularly in the Abundance model (green) (Figure 2G, top row; Supplemental Figures 2A,C). This is consistent with the idea that tetraploidy serves as an intermediate enabling a near-triploid karyotype that is common in many cancers (Bielski et al., 2018; Lopez et al., 2020). With regard to MKV, diploid populations overall, and the Driver model in particular, tend to exhibit increased MKV in fewer steps, suggesting a more rapid increase in diversity (Figure 2G, bottom row). This is explained by the model assumption that a maximum of one of any particular chromosome mis-segregates per division, and one mis-segregation is a greater fraction relative to diploidy than triploidy. Taken together, these models recapitulate key aspects of prior characteristics of aneuploidy and ploidy status on tumor evolution, lending credence to our model. Further, they illustrate that it may not be possible to directly infer mis-segregation rates, a measure of CIN, solely by sampling karyotypes in a human tumor without accounting for selection. However, this model serves as a framework for quantitative inference of mis-segregation rates form the diversity in tumor-derived single-cell DNA sequencing.

### Karyotype diversity depends profoundly on selection modality

Some current measures of CIN are derived from the aneuploidy diversity in the population. Yet, our model suggests that selection may profoundly shape the karyotype variance in a population. To evaluate this further, we considered the case where CIN is a property of an incipient tumor, but then is turned off (P_misseg_ = 0) after 200 steps (Figure 3A). Over an additional 800 steps of selection, tumors could retain high aneuploidy (top arrow), revert to a small number of distinct clones (middle arrow), or return to a near-diploid state (bottom arrow). For 200 steps, we simulated the karyotypic evolution of populations at low rates of chromosome mis-segregation, akin to what has been observed microscopically (P_misseg_ = 0.022 per chromosome; ∼ 1 chromosome per division) (Bakhoum et al., 2009; Weaver et al., 2007), and continued selection for the remaining timescale (1000 steps) with moderate selective pressure (S=5). This early CIN mimics the chromosomal instability in early tumor evolution as reported to occur in breast cancer (Gao et al., 2016) (Figure 3A).

**Figure 3.**
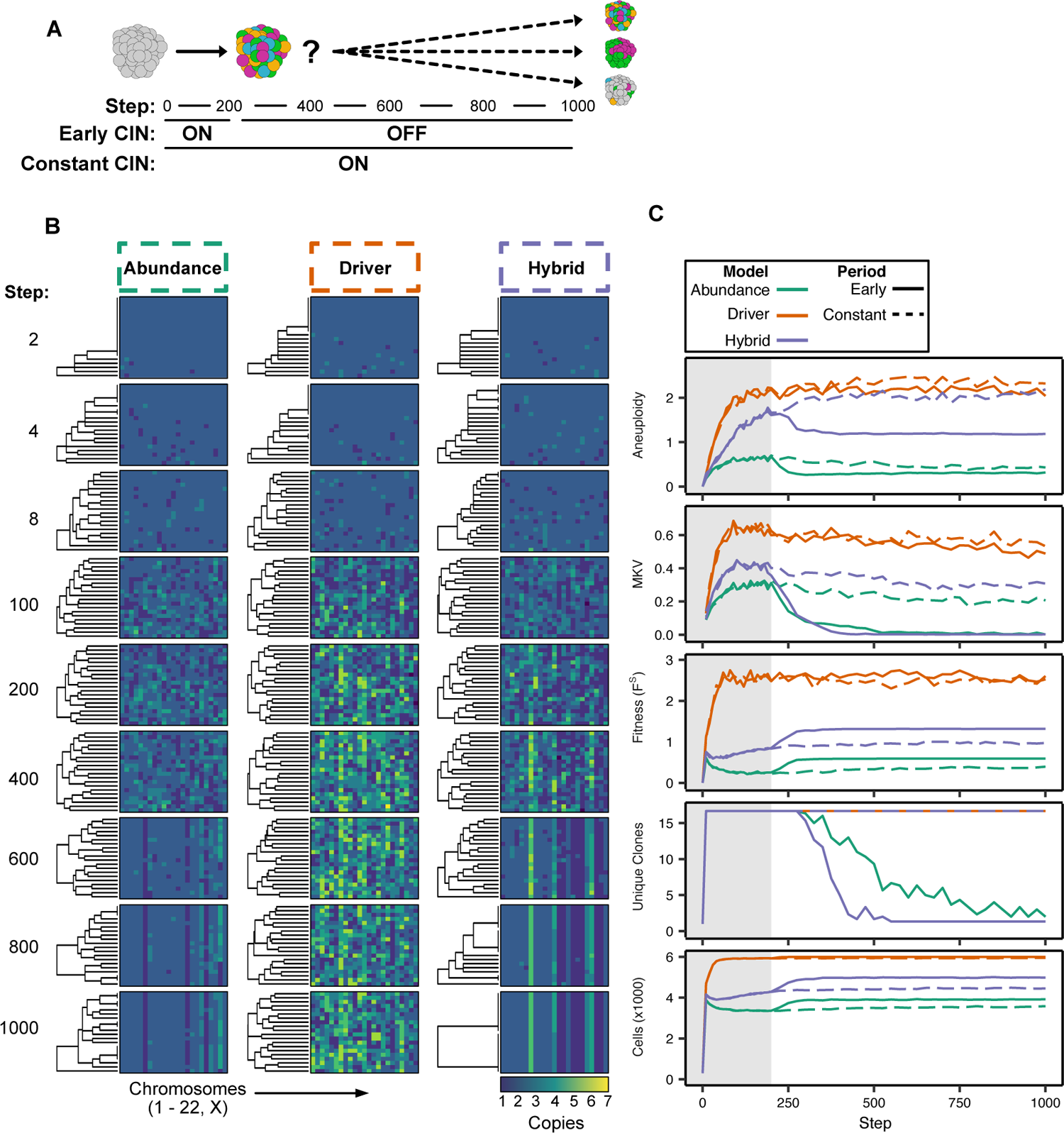
Karyotype diversity depends profoundly on selection modality. (A) Scheme for simulation experiment to observe the emergence of unique clones. Early CIN simulations were run with CIN for 200 steps and a further 800 without. (B) Heatmaps depicting the chromosome copy number profiles of a subset (n = 20 out of 300 sampled cells) of the simulated population with early CIN over time under each model of karyotype selection. Pmisseg = 0.022; S = 5. (C) Time lapse analysis of a subset (n=20 out of 300) of the simulated population. Grey box depicts time with CIN. Aneuploidy is measured as the mean of the variances taken within each cell’s karyotype (Supplemental Figure 5A). Unique clones are any cell with a unique karyotype. Lines depict the mean of n=3 simulations.

We visualized the karyotype profiles of individual cells as heatmaps over time (Figure 3B). Under the Abundance selection model (left), the cell population indeed diversified over 200 steps, but then became more uniform, with selection markedly reducing the number of unique clones and, hence, karyotypic variance. The population generated but did not maintain a high degree of aneuploidy (mean intra-karyotype variance; Supplemental Figure 5A) as a result of decreased fitness over the entire population. By contrast, the Driver Density model (middle) maintained karyotypic diversity at later time steps. Accordingly, the number of unique clones did not decrease over time. This indicates that the model does not strongly select against the karyotypically distinct clones that have F^S^>2, thus a P_division_ = 1 (where cells will divide at every time step). The Hybrid model (right) exhibited an intermediate level of aneuploidy and MKV generated during the first 200 steps; after which, many, but not all clones were removed over time (Figures 3B,C). These data indicate that the development and maintenance of karyotype diversity over time depends profoundly on the modality of karyotype selection.

### Topological features of simulated phylogenies delineate CIN rate and karyotype selection

Given a model capable of recapitulating diversity and selective pressures, how is it possible to account for selection to infer P_misseg_ as a measure of CIN from an observed population of cells? Phylogenetic trees are a useful tool to infer evolutionary processes of genetic diversification and selection. Moreover, the topology of the phylogenetic tree has been used as a quantitative measure of the underlying evolutionary processes they result from (Colijn and Plazzotta, 2018; Dayarian and Shraiman, 2014; Manceau et al., 2015; Neher et al., 2014; Scott et al., 2019).

Here, chromosome mis-segregation gives rise to variable levels of karyotypic heterogeneity within a population, which is then shaped—and confounded—by selection. We sought to understand how we can utilize chromosome copy number-based phylogenic reconstruction to disambiguate these factors by quantifying the topological features of these simulated phylogenetic trees. These include discrete features such as ‘cherries’—two tips that share a direct ancestor—and ‘pitchforks—a clade with three tips. Additionally, we considered a broader metric of topology, the Colless index, which measures the imbalance or asymmetry of the entire tree (Figure 4A). To understand how these measures are affected by selection in our model, we looked explicitly at the phylogenetic reconstructions of 300 random cells from each simulation performed over a full range of selective pressures taken at 60 time steps (∼30 divisions)(Figure 4B). As seen previously, aneuploidy and MKV decreased with selective pressure in a trend that becomes more robust as mis-segregation rate increases. As expected, the Colless indices for these simulations appear to increase with selective pressure, which tends to generate asymmetry through selection on early aneuploid karyotypes (Figure 4C). Accordingly, this imbalance is apparent in phylogenetic reconstructions of simulated populations (Figure 4D). Cherries and pitchforks also vary, though less consistently. Yet cherries tend to diminish with very high selective pressure, probably due to selection against variation in aneuploidies even in the latest generations prior to sampling (Figure 4C, 4^th^ row).

**Figure 4.**
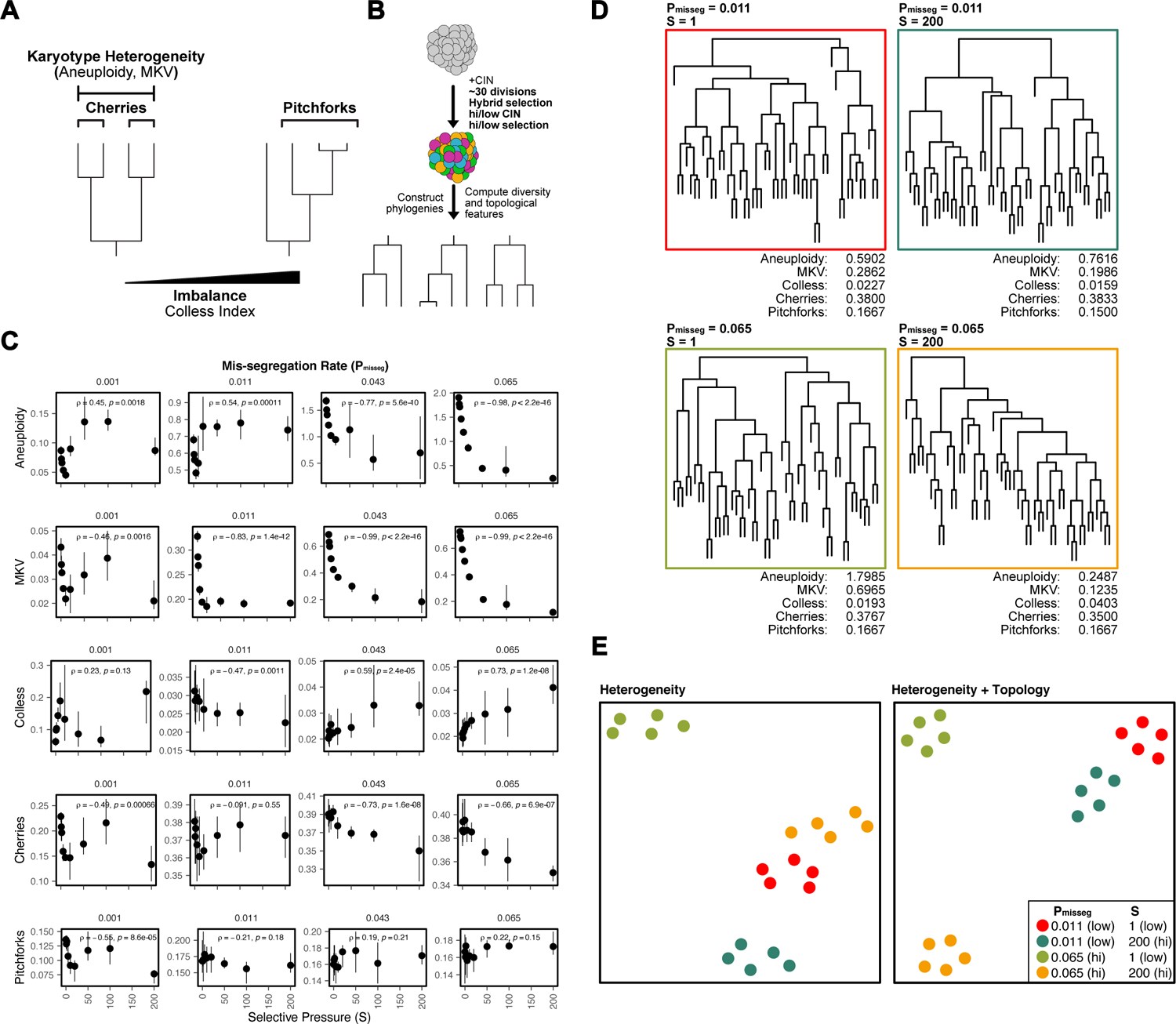
Topological features of simulated phylogenies delineate CIN rate and karyotype selection. (A) Quantifiable features of karyotypically diverse populations. Heterogeneity between and within karyotypes is described by MKV and aneuploidy (inter- and intra-karyotype variance, see Materials & Methods). We also quantify discrete topological features of phylogenetic trees, such as cherries (tip pairs) and pitchforks (3-tip groups), and a whole-tree measure of imbalance (or asymmetry), the Colless index. (B) Scheme to test how CIN and selection influence the phylogenetic topology of simulated populations. (C) Computed heterogeneity (aneuploidy and MKV) and topology (Colless index, cherries, pitchforks) summary statistics under varying Pmisseg and S values. MKV is normalized to the average ploidy of the population. Topological measures are normalized to population size. Spearman rank correlation coefficients and p-values are displayed (n = 5 simulations). (D) Representative phylogenies for each hi/low CIN, hi/low selection parameter combination and their computed summary statistics. Each phylogeny represents n = 50 out of 300 cells for each simulation. (E) Dimensionality reduction of all simulations for each hi/low CIN, hi/low selection parameter combination using measures of karyotype heterogeneity only (left; MKV and aneuploidy) or measures of karyotype heterogeneity and phylogenetic topology (right; MKV, aneuploidy, Colless index, cherries, and pitchforks).

To characterize how well these measures retain information about the simulation parameters, we performed dimensionality reduction with measures of karyotype heterogeneity alone (MKV and aneuploidy) and with both measures of karyotype heterogeneity and phylogenetic topology (Figure 4E). This analysis indicates that when considering heterogeneity alone simulations performed under high CIN/high selection (yellow) and low CIN/low selection (red) associate closely, meaning these measures of heterogeneity are not sufficient to distinguish these disparate conditions (Figure 4E, left). These similarities arise because high selection can mask the heterogeneity expected from high CIN. By contrast, combining measures of heterogeneity with those of phylogenetic topology can better discriminate between simulations with disparate levels of CIN and selection (Figure 4E, right). This provides further evidence that measures of heterogeneity alone are not sufficient to infer CIN due to the confounding effects of selection, particularly when the nature of selection is unclear or can vary. Together these results indicate that phylogenetic topology is able to delineate the levels of selective pressure and rates of chromosome mis-segregation under which these simulations were performed. Further, they indicate that considering phylogenetic topology in single-cell populations may be a suitable method for correcting for selective pressure when estimating the rate of chromosome mis-segregation from measures of karyotype diversity.

### Experimental chromosome mis-segregation measured by Bayesian inference

To experimentally validate quantitative measures of CIN, we generated a high rate of chromosome mis-segregation with a clinically relevant concentration of paclitaxel (Taxol) over 48 hours (Figure 5A). We treated CAL51 breast cancer cells with either a DMSO control or 20 nM paclitaxel, which is expected to generate widespread aneuploidy due to chromosome mis-segregation on multipolar mitotic spindles (Zasadil et al., 2014), which we verified in this experiment (Supplemental Figure 4A). At this short timescale we assume cells have undergone only 1-2 mitoses and we observe broadened DNA content distributions by flow cytometry (Supplemental Figure 4B). Using low-coverage scDNAseq data, we characterized the karyotypes of 38 DMSO- and 134 paclitaxel-treated cells. As expected, we observe a high penetrance and extensive degree of aneuploidy generated via paclitaxel, which contrasts with low variance in the control (Figure 5B). Additionally, the mean of the resultant aneuploid karyotypes for each chromosome still resembled those of bulk-sequenced cells, highlighting that bulk-sequencing is an ensemble average, and does not detect the true variation in the population, particularly with balanced mis-segregation events (Figure 5B, single-cell mean and bulk). In quantifying the absolute deviation from the modal control karyotype in each cell, and assuming a single mitosis, we found that cells that undergo mitosis in the presence of 20 nM paclitaxel mis-segregate 18.5 ± 0.5—a P_misseg_ of ∼0.42 (Figure 5C). Moreover, we observe a close association between the incidence of nullisomy observed experimentally to simulations carried out under this mis-segregation rate over 1-2 divisions (Figure 5D).

**Figure 5.**
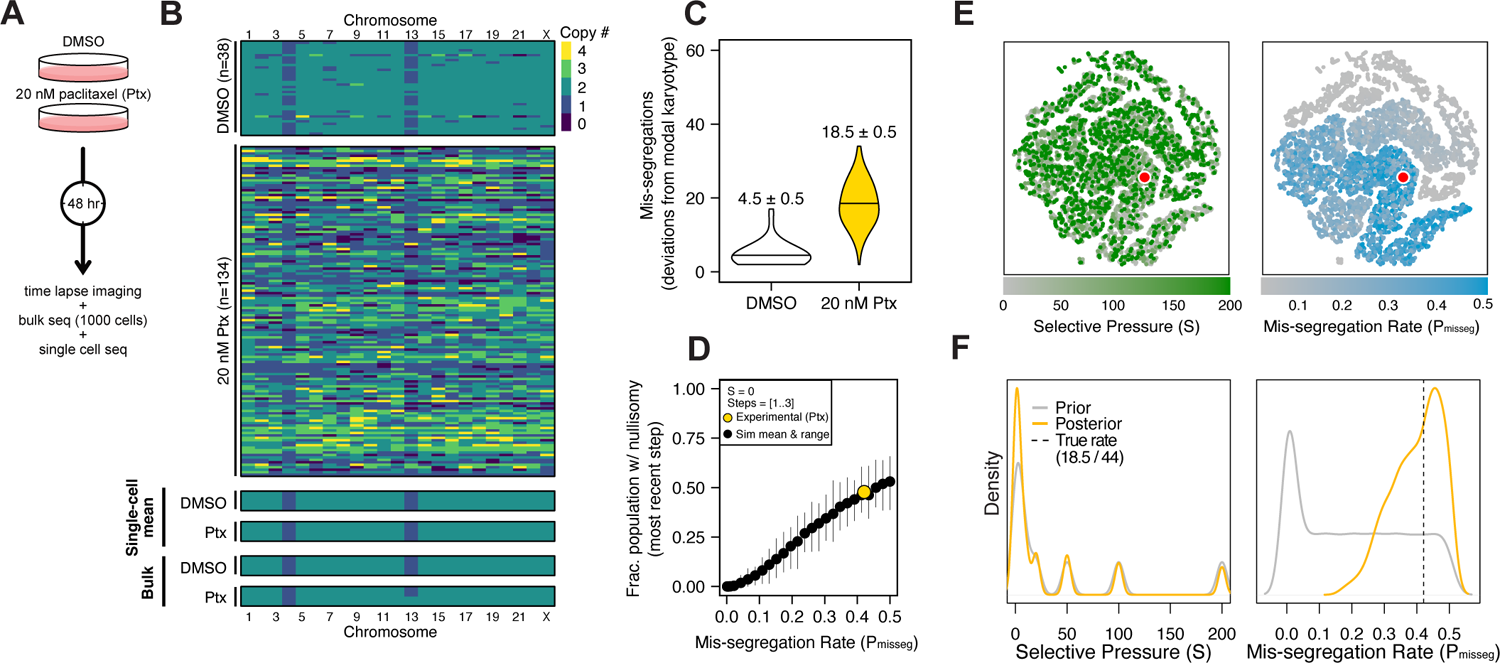
Experimental chromosome mis-segregation measured by Bayesian inference. (A) Experimental scheme. Cal51 cells were treated with either DMSO or 20 nM paclitaxel for 48 hours prior to further analysis by time lapse imaging, bulk DNA sequencing, and scDNAseq. (B) Heatmaps showing copy number profiles derived from scDNAseq data, single cell copy number averages, and bulk DNA sequencing. (C) Observed mis-segregations calculated as the absolute sum of deviations from the observed modal karyotype of the control. (D) Observed incidence of nullisomy (fraction∼0.5) in paclitaxel-treated cells plotted against the observed mis-segregation rate (Pmisseg,true = 18.5/44 = 0.42) overlaid on simulated data from the first 3 time steps (∼1.5 generations) under the Hybrid model. (E) Dimensionality reduction analysis of population summary statistics (aneuploidy, MKV, Colless index, cherries, and pitchforks) from the first 3 time steps of all simulations performed under the Hybrid model. (F) Prior (grey) and posterior (gold) distributions from Approximate Bayesian computation analysis using population summary statistics computed from the paclitaxel-treated cells. Only the first 3 time steps of simulation data were used. Dashed line represents the experimentally observed mis-segregation rate.

In this instance, while we were able to estimate mis-segregation rate by calculating absolute deviation from the modal karyotype after a single aberrant cell division. However, such an analysis would be inappropriate for long term experiments, or real tumors, where new aneuploid cells may be subject to selection. Accordingly, we sought to infer the parameters of this experiment—the mis-segregation rate of 18.5 chromosomes per division and low selection— using only measures of aneuploidy, variance, and phylogenetic topology as shown previously. To display this, we used dimensionality reduction to ensure that observed measures from the paclitaxel-treated Cal51 population fell within the space of those observed from simulated populations over 1-3 steps (0.5-1.5 generations). Indeed, the experimental data mapped to those from simulations using high mis-segregation rates and simulated selection pressures were poorly separated, indicating karyotype selection is not a major factor within this time frame (red point, Figure 5E). However, this comparison does not provide a quantitative measure of CIN. Instead, parameter inference via approximate Bayesian computation (ABC) is well suited for this purpose.

By providing these same metrics derived from simulated populations evolved under wide-ranging, uniform distributions of evolutionary parameters (a prior distribution), ABC can determine the most likely evolutionary parameters that produce the observed pattern without biased *a priori* estimations (the posterior distribution). We utilized an ABC framework (Csillery et al., 2012) on our simulated dataset to infer the chromosome mis-segregation rate and selective pressure observed in the paclitaxel-treated cells (see Materials and Methods). We then used the experimentally observed mis-segregation rate as a benchmark to optimally select a panel of measures for parameter inference (Supplemental Figure 5) and selected the following five metrics to use concurrently in our ABC analysis: mean aneuploidy, MKV, the Colless index (a phylogenetic balance index) and the population size-normalized number of cherries and pitchforks (discrete topological features of phylogenetic trees). In doing so, this analysis inferred a chromosome mis-segregation rate of 17.5±0.14 chromosomes (mean ± SE), an estimate that compares favorably with the experimentally observed rate of 18.5±0.5. The range and density of accepted values for selective pressure spanned the entire posterior distribution, meaning that selection had little bearing on the result, regardless of the magnitude, at this time point (Figure 5F). This is consistent with the absence of selection in a 48-hour experiment. In short, this experimental case allowed us to validate ABC-derived mis-segregation rate as a measure of CIN, using an experimentally determined mis-segregation rate as confirmation. Importantly, prior estimations of mis-segregation rate selective pressure were not required to develop this quantitative measure of CIN.

Together, these data indicate that combining simulated and observed metrics of population diversity and structure with a Bayesian framework for parameter inference may be a flexible method of quantifying the evolutionary forces associated with CIN. Moreover, using this method has revealed the hitherto unreported potential extent of chromosome mis-segregation induced by a clinically relevant concentration of the successful chemotherapeutic paclitaxel.

### Minimum sampling of karyotype heterogeneity

The cost of high-throughput DNA sequencing of single cells is often cited as a limitation to clinical implementation. In part, the cost can be limited by low-coverage sequencing which is sufficient to estimate the density of reads across the genome. Further, it may be possible to minimize the number of cells that are sampled to get a robust estimate of CIN, though sampling too few cells may result in inaccurate measurements. Accordingly, we sought to understand how sampling impacts measurement of mis-segregation rates using approximate Bayesian computation. We first took 5 random samples from the population of paclitaxel-treated cells each at various sample sizes (Supplemental Figure 6A). We then inferred the mis-segregation rate in each sample and identified the sample size that surpasses an average of 90% accuracy and a low standard error of measurement. We found the low sample sizes (n=[10..40]) suffer underestimation of the known mis-segregation rate in paclitaxel-treated cells (Supplemental Figure 6B). While the mean percent accuracy levels off above 90% at about 50 cells (Supplemental Figure 6C), the standard error of measurements stabilizes around 70 cells (Supplemental Figure 6D). Random samples of 70 paclitaxel-treated cells had a percent accuracy of 95.1% and a standard error of 0.03 (±1.3 chromosomes per division). We repeated this analysis using simulated data from the Hybrid selection model and a range of mis-segregation rates spanning what is observed in cancer and non-cancer cultures (P_misseg_ ≤ 0.022; Figure 6E). We again found a range of sample sizes whose inferred mis-segregation rates underestimate the known value from those simulations (n=[20..160]; Supplemental Figures 6E,F). Across all mis-segregation rates and selective pressures, random samples of 180 cells had a median percent accuracy of 92% and median standard error of 0.0002 (± 0.0092 chromosomes per division). The difference in optimal sample sizes between the paclitaxel-treated population and the simulated population is notable and likely due to the presence of ‘clonal’ structures in the simulated population. While the paclitaxel treatment resulted in a uniformly high degree of aneuploidy and little evidence of karyotype selection, the simulated populations after 60 steps (∼30 generations) have discrete copy number clusters that may not be captured in each random sample. To verify this, we repeated the analysis using only data from the first time step, prior to the onset of karyotype selection (Supplemental Figure 6H). In this case, we found that the sample size needed to achieve a median 90% accuracy over all simulations in this context is 100 cells, at which point the standard error for P_misseg_ is 0.0068 (placing measures within ± 0.31 chromosomes per division; Supplemental Figures 6I,J). Thus, a larger number of cells is required in the context of long-term karyotype selection than a more acute time scale, such as we see with paclitaxel.

**Figure 6.**
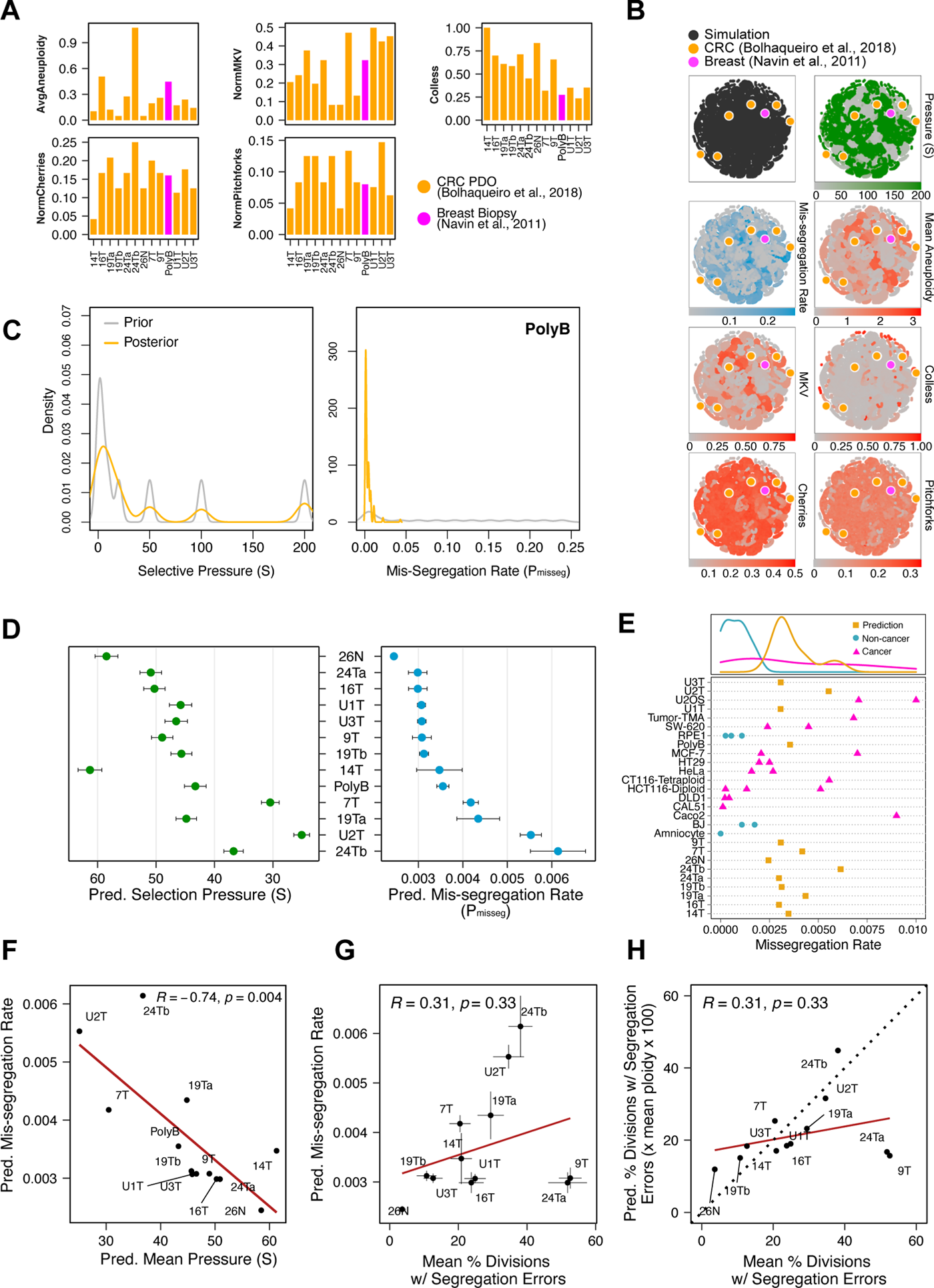
Inferring chromosome mis-segregation rates in tumors and organoids. (A) Computed population summary statistics for colorectal cancer (CRC) patient-derived organoids (PDOs) and breast biopsy scDNAseq datasets from Bolhaqueiro et al., (2019) (gold) and Navin et al., (2011) (pink). (B) Dimensionality reduction analysis of population summary statistics showing biological observations overlaid on, and found within, the space of simulated observations. Point colors show the distributions of simulation parameters and summary statistics for all simulations with Pmisseg ≤ 0.3. (C) Plots showing the prior (grey) and posterior (gold) distributions of Pmisseg and S values from the approximate Bayesian computation analysis of the breast biopsy sample from Navin et al., 2011. ABC analyses were performed using a tolerance threshold of 0.05 to reject dissimilar simulated observations (see Materials and Methods). Only simulations with Pmisseg ≤ 0.3 were used for parameter inference. D) Inferred selective pressures and mis-segregation rates from each scDNAseq dataset (mean and SEM of accepted values). (E) Pearson correlation of predicted mis-segregation rates and predicted selective pressures in CRC PDOs. (F) Pearson correlation of predicted mis-segregation rates and the incidence of observed segregation errors in CRC PDOs from Bolhaqueiro et al., 2019. Error bars represent SEM values. (G) Pearson correlation of observed incidence of segregation errors to ploidy-corrected prediction of the observed incidence of segregation errors. These values assume the involvement of 1 chromosome per observed error and are calculated as the (predicted mis-segregation rate) x (mean number of chromosomes observed per cell) x 100. Dotted line = 1:1 reference. (H) Predicted mis-segregation rates in CRC PDOs and a breast biopsy plotted with approximated mis-segregation rates observed in cancer (blue triangle) and non-cancer (red circle) models (primarily cell lines) from previous studies (Supplemental Table 2; see Materials and Methods). The predicted mis-segregation rates in these cancer-derived samples fall within those observed in cancer cell lines and above those of non-cancer cell lines.

In conclusion, we recommend using 180 cells from a single sampled site which, at biologically relevant time scales and rates of mis-segregation, provides ≥90% accuracy. These data represent, to our knowledge, the first analysis of how sample size for single-cell sequencing affects the accuracy and measurement of chromosome mis-segregation rates.

### Inferring chromosome mis-segregation rates in tumors and organoids

To determine if this framework is clinically applicable, we sought out previously published scDNAseq datasets derived from tumor samples and patient-derived organoids (PDO) (Bolhaqueiro et al., 2019; Navin et al., 2011). Importantly, the data from Bolhaqueiro et al. include sample-matched live cell imaging data in colorectal cancer PDOs, with direct observation of chromosome mis-segregation events to compare with inferred measures. We established our panel of measurements on these populations (Figure 6A) then constrained our prior distribution of simulated data to those with chromosome mis-segregation rates below 0.25, because above this threshold populations were unstable and typically died off after ∼20 time steps (Figure 2C). We then confirmed these datasets were within the space of simulation data from the Hybrid model (Figure 6B). Next, we performed ABC analysis on these datasets to infer rates of chromosome mis-segregation and levels of selection pressure (Figure 6C; Supplemental Figure 7). Figure 6C illustrates the results for the biopsied breast tumor from Navin et al., 2011, illustrating the distribution of parameters used for simulations that gave the most similar results. When applying ABC to infer paramters for various samples (Figure 6D), we find these fall within rates of mis-segregation of about 0.001 to 0.007. Applied to a near-diploid cell, this would translate to a range of about 5-36% of cell divisions having one chromosome mis-segregation. Importantly, these putative rates of chromosome mis-segregation fall within the range of approximated *per chromosome* rates experimentally observed in cancer cell lines and human tumors (Bakhoum et al., 2014, 2011, 2009; Dewhurst et al., 2014; Nicholson et al., 2015; Orr et al., 2016; Thompson and Compton, 2008; Worrall et al., 2018; Zasadil et al., 2014). Higher inferred mis-segregation rates significantly correlated with lower inferred selection experienced in these samples (Figure 6F). Notable examples are the sample from normal colon tissue (26N) and a near-triploid sample (24Tb), which were inferred to be experiencing one of the highest and lowest levels of selection respectively. This is consistent with the idea that normal tissue does not well tolerate aneuploidy whereas high-ploidy tumors better tolerate CIN (Dewhurst et al., 2014; Knouse et al., 2014; Lopez et al., 2020; Pfau et al., 2016). Additionally, by estimating the number of generations experienced by these populations, we found that the breast biopsy sample spent more time diversifying prior to sampling than the CRC PDOs (Supplemental Figures 8A,B). This seems appropriate as the CRC PDOs began as single cell clones with relatively little time in culture. However, these values were estimated using a post-hoc calculation of the average P_div_ of each population. This is only accurate insofar as the chromosome scores for each model truly represent aneuploid fitness in these samples but may serve as a relative time scale for each sample. As further validation, we compared these inferred mis-segregation rates from CRC PDOs with those directly measured in live imaging. There was a strong correlation but for two outliers—9T and 24Ta (Figure 6G). In fact, when adjusting to the same scale and correcting for cell ploidy, these data follow a strong positive linear trend, excepting the outliers (Figure 6H). Overall, these results indicate that the inferred measures using approximate Bayesian computation and scDNAseq account for selection and provide a quantitative measure of CIN.

## DISCUSSION

The clinical assessment of mutations, short indels, and microsatellite instability in human cancer determined by short-read sequencing currently guide clinical care. By contrast, CIN is highly prevalent, yet has remained largely intractable to clinical measures. Single-cell DNA sequencing now promises detailed karyotypic analysis across hundreds of cells, yet selective pressure is expected to suppress the observed karyotype heterogeneity within a tumor. We therefor conclude that optimal clinical measures of CIN require an approach that employs scDNAseq data, yet account for selective pressure to infer the underlying rates of chromosome mis-segregation events.

Despite the existing major limitations with quantitative measures of CIN, emerging evidence hits at its utility as a biomarker to predict benefit to cancer therapy. For example, preliminary data suggests that CIN measures can predict therapeutic response to paclitaxel (Janssen et al., 2009; Swanton et al., 2009). However, these have not been implemented in the clinic, in part because existing measures of CIN have had significant limitations. FISH and histological analysis of mitotic abnormalities are limited in quantifying specific chromosomes or requiring highly proliferative tumor types, such as lymphomas and leukemia. Gene expression profiles have been proposed to correlate with CIN among populations of tumor samples (Carter et al., 2006), though they happen to correlate better with tumor proliferation (Sheltzer, 2014); in any case, they are correlations across populations of tumors, not suitable as an individualized diagnostic. Computational modeling has been used to explore evolution in the context of numerical CIN and karyotype selection (Elizalde et al., 2018; Gao et al., 2016; Gusev et al., 2001, 2000; Laughney et al., 2015). However, no prior study, to our knowledge, has developed quantitative approaches to measure CIN while accounting for cellular selection.

Previous studies using single-cell sequencing identified surprisingly low karyotypic variance in human tumors including breast cancer (Gao et al., 2016; Kim et al., 2018; Wang et al., 2014) and colorectal and ovarian cancer organoids (Bolhaqueiro et al., 2019; Nelson et al., 2020). It has been difficult to understand these findings in the light of widespread CIN in human cancer (Sheltzer and Amon, 2011; Silk et al., 2013; Vasudevan et al., 2020; Weaver et al., 2007; Weaver and Cleveland, 2009). The best explanation of this apparent paradox is selection, which moderates karyotypic variance. Accounting for this, we are able to infer rates of chromosome mis-segregation in tumors or PDOs well within the range of rates observed microscopically in cancer cell lines. Additionally, no previous work, to our knowledge, has estimated the required sample size to infer CIN from scDNAseq data.

As described by others (Dewhurst et al., 2014; Lopez et al., 2020), and consistent with our findings, early emergence of polyploid cells can markedly reduce apparent selection, leading to an elevated karyotype diversity over time. While we do not explicitly induce whole genome doubling (WGD) in these simulations, populations that begin either diploid or tetraploid converge on near-triploid karyotypes over time, supporting the notion that WGD can act as an evolutionary bridge to highly aneuploid karyotypes. Notably, our analysis indicates the only sample with apparent polyploidy experienced among the lowest levels of karyotype selection.

In some early studies, CIN is considered a binary process—present or absent. We assumed that CIN measures are scalar, not binary, and measure this by rate of chromosome mis-segregation per division. A scalar is appropriate if for example there was a consistent probability of chromosome mis segregation per division. However, we recognize that some mechanisms may not well adhere to this simplified model of CIN. For example, tumors with centrosome amplification undergo bipolar division without mis-segregation, or a multipolar division with extensive mis-segregation. Another possibility is that CIN could result in the misregulation of genes that further modify the rate of CIN. Our model does not yet account for punctuated behavior or changing rates of CIN.

Another limitation of our model and inference of mis-segregation rates is the possibility of structural chromosomal instability—some mechanisms of instability such as breakage-fusion-breakage can result in structural aberrations that may differ cell-by-cell and result in the formation of ‘pseudo-‘ or ‘derivative-chromosomes’ through *trans*-chromosomal rearrangement. Our current model is limited to whole-chromosome measures. Additional measures, such as homologous recombination deficiency (HRD) scores, which correlate with various measures of genomic instability (Marquard et al., 2015), would be better suited to quantifying structural variation and could feasibly be integrated with this framework in a more genetically granular iteration. However, we note that our approach could be used to quantify CIN even in the setting of structural variation.

A final limitation of our approach is we used previous estimates of cellular selection in our agent-based model and used these selection models to infer quantitative measures of CIN. While this approach seems to perform well in estimates of mis-segregation rates, we recognize that the selection models do not necessarily represent the real selective pressures on distinct aneuploidies. Future investigations are necessary to directly measure the selective pressure of distinct aneuploidies—a project that is now within technological reach. It is also a possibility that the selective pressures could be influenced by cell type (Auslander et al., 2019; Dürrbaum et al., 2014; Sack et al., 2018; Starostik et al., 2020), tumor cell genetics (Foijer et al., 2014; Grim et al., 2012; López-García et al., 2017; Simoes-Sousa et al., 2018; Soto et al., 2017), and the microenvironment (Hoevenaar et al., 2020).

In summation, we developed a theoretical and experimental framework for quantitative measure of chromosomal instability in human cancer. This framework accounts for selective pressure within tumors and employs Approximate Bayesian Computation, a commonly used analysis in evolutionary biology. Additionally, we determined that low-coverage single-cell DNA sequencing of at least 180 cells from a human tumor sample is sufficient to get an accurate (>90% accuracy) and reproducible measure of CIN. In conclusion, this work sets the stage for standardized quantitative measures of CIN that promise to clarify the underlying causes, consequences, and clinical utility of this nearly universal form of genomic instability.

## METHODS

### Underlying assumptions for models of CIN and selective pressure

1. Chromosome mis-segregation rate is defined as the number of chromosome mis-segregation events that occur per cellular division.
2. Cell division always results in 2 daughter cells.
3. P_misseg,c_ is assigned uniformly for each cell in a population and for each chromosome.
4. Cells die when the copy number of any chromosome is equal to 0 or exceeding 6.
5. Steps are based on the rate of division of euploid cells. We assume a probability of division (P_division_) of 0.5, or half of the population divides every step, for euploid populations. This probabilistic division is to mimic the asynchrony of cellular proliferation and to allow for positive selection, where some cells may divide more rapidly than their euploid ancestors.
6. No chromosome is more likely to mis-segregate than any other.
7. Each parental chromosome may only be mis-segregated once per division. For example, for a tetraploid cell, the two maternal chromosomes are never simultaneously mis-segregated.

### Modeling chromosome mis-segregation

In each model, numerical scores are assigned to each chromosome, the sum of which represents the fitness of the karyotype (Figure 1B). Founder populations of 100 individual ‘agents’ or cells are grown probabilistically over 60 time steps where each cells probability of mitosis (P_mitosis_) is equal one half its fitness score. For example, a euploid cell with perfect fitness will undergo mitosis every other time step. Chromosomes are assigned fitness contribution (f_c_) scores based on their estimated gene abundance (normalized to the estimated total number of genes) or driver density (normalized to the sum of all chromosome scores) In this way, chromosomes with higher baseline fitness values contribute more to cellular fitness (Figure 1B). Cells are then passed through an iterative framework as follows:

1. Chromosomes’ f_c_ values are modified to generate a Contextual Fitness Score (CFS) according to the model used. In the Gene Abundance model (Figure 1, upper-left), f_c_ values are modified such that these values decrease if the copy number of that chromosome deviates from the average ploidy of the population. In the model of TOE densities (Figure 1, lower-left), the CFS is only dependent on the copy number of that chromosome, irrespective of ploidy. In the hybrid model, these two processes are weighted equally and averaged. With these chromosome scores assigned, the fitness of each cell in the starting population is equal to one (F = 1) as each starting cell is euploid
2. A cell’s probability of entering mitosis is dependent on its fitness (F) which is the sum of all CFS values.
3. Fitness can be modified (F_M_) by applying selective pressure (F_M_ = F^S^).
4. If a cell has a copy number of 0 or greater than 6 for any chromosome as a result of the most recent mitosis, the cell is killed. Additionally, to reduce computational constraints, populations are limited to ∼4500 cells using a process wherein, when this limit is reached, a random half of all cells are deleted (Figure 2B). This process does not impact the results of long-term experiments as each cell uniformly has the same probability of being removed and the same probability of experiencing a chromosome mis-segregation event (Supplemental Figure 3).
5. Cells undergo mitosis probabilistically based on their F_M_ values. Cells with larger F_M_ values will divide more rapidly.
6. If a cell enters mitosis, chromosomes are mis-segregated probabilistically by choosing a random integer (*R*) from a uniform distribution of length *L* and determining if this number is less than 100. This is done for each parental chromosome such that P_misseg,c_ = P(*R* < 100) where *L* is predetermined.
7. The model stops when 60 time steps have been reached.

Models were coded in the agent-based platform, NetLogo 6.0.4 (Wilensky, 1999). In general, a euploid population of cells (agents) are initialized in space with integer values assigned to each chromosome, with respect to ‘maternal’ and ‘paternal’ chromosomes (i.e. for diploid cells, chromosome copy number states = 1). At each step, the fitness (F) is modified (F_M_) by the selective pressure (S) such that (F_M_ = F^S^). Thus, when S = 0, F_M_ = 1.

Each cell probabilistically determines if it undergoes mitosis based on its fitness value. A random number is drawn from *R* ∼ *U*([0..200]) and cells are made to divide when *R* ≤ F_M_ x 100 such that P(mitosis) = P(*R* ≤ F_M_ x 100) = F_M_ / 2.

If a cell is determined to divide, chromosomes are similarly probabilistically determined to be retained or mis-segregated. Where *L* is defined by the pre-selected probability of chromosome mis-segregation, *R* ∼ *U*([0..*L*]) is assessed for each chromosome (C). Thus, chromosomes are mis-segregated when P_misseg,c_ = P(*R* < 100). For example, when *L* = 4600, P_misseg,c_ = 1/46 and a single chromosome from a cell’s karyotype will be mis-segregated on average. The probability that a cell gains or loses a chromosome are equal (P=0.5). When mis-segregation is completed, a copy of that cell will be generated with the opposite result of the previous mis-segregations.

At this point, cells with any chromosome copy number (n_c_) equaling 0 or exceeding 6 are deleted and remaining cells recalculate their fitness values under the selected selection modality. The simulation ends when the number of time steps reaches the pre-selected timeframe. Populations were limited to ∼4500 cells by randomly removing half of the population at each time step once this number is reached.

### Modeling karyotypic selection modalities

At each simulation time step, fitness is re-calculated based on their updated karyotype. These fitness values determine if they undergo mitosis in the next round. However, the modality of selection changes how those karyotypes are assessed. We implement 3 separate selection modalities based on the chromosomal features of 1) gene abundance, 2) driver density, or 3) a hybrid of both. The scores that are generated in each produce a fitness value (F) that can then be subjected to pressure (S) as described above.

## 1. Selection on Gene Abundance

The Gene Abundance modality relies on the concept of gene dosage stoichiometry where the aneuploid karyotypes are selected against and that the extent of negative selection scales with the severity of aneuploidy and the identity and gene abundance on the aneuploid chromosomes (Sheltzer and Amon, 2011). Gene abundance values for each chromosome were retrieved from GRCh38.p13 (https://www.ncbi.nlm.nih.gov/genome/51). Chromosome fitness contribution scores (f_c_) were generated by normalizing their gene abundance to the total number of genes in the genome. Thus, the sum of all based normalized gene abundance scores is 1. These base values are then modified under the gene abundance model to generate a contextual fitness score (CFS_GA,c_) at each time step such that…

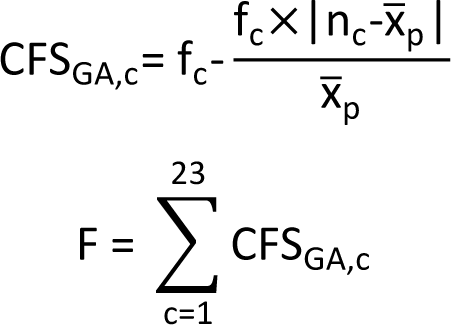

… where x#_p_ is the average ploidy of the population and n_c_ is the chromosome copy number. This modality then dictates that the fitness contribution of a chromosome declines as its distance from the average ploidy increases and that the magnitude of this effect is dependent on the size of the chromosome.

## 2. Selection on Driver Densities

The Driver Density modality relies on assigned fitness values to chromosomes based on their relative density of tumor suppressor genes, essential genes, and oncogenes. These values were derived from a pan-cancer analysis (TCGA) of the frequency of mutation of these genes and their location in the genome (Davoli et al., 2013). These so-called ‘chrom-scores’ correlate with the frequency with which chromosomes are found to be amplified in the genome. Thus, this selection modality benefits cells that have maximized the density of oncogenes and essential genes to tumor suppressors through chromosome mis-segregation. Chrom-scores were normalized to the sum of their values and assigned to their respective chromosomes. Chromosome X did not receive a chrom-score, thus its assigned score was 0. These base scores (TOE_c_)were then modified to generate a contextual fitness score such that…

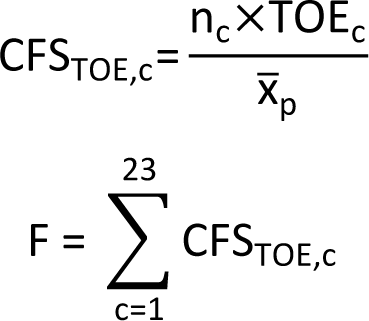

## 3. Selection on Gene Abundance and Driver Densities (Hybrid)

The hybrid model relies on selection on both gene abundance and driver densities. CFS_TOE,c_ and CFS_GA,c_ are both calculated and averaged such that…

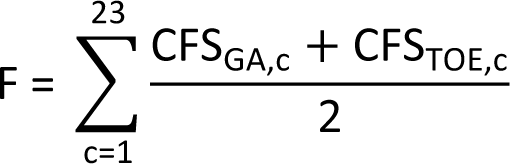

### Analysis of population diversity and topological features of phylogenetic trees

Phylogenetic trees were reconstructed from chromosome copy number profiles from live and simulated cells by calculating pairwise Euclidean distance matrices and performing complete-linkage clustering in R (R Core Team, 2013). Phylogenetic tree topology measurements were performed in R using the package phyloTop v2.1.1 (Kendall et al., 2018). Sackin and Colless indices of tree imbalance were calculated, normalizing to the number of tree tips. Cherry and pitchfork number were also normalized to the size of the tree. MKV is taken as the variance of individual chromosomes taken across the population, averaged across all chromosomes, then normalized to the average ploidy of the population. Average aneuploidy is calculated as the variance within a single cell’s karyotype averaged across the population.

### Approximate Bayesian computation

Approximate Bayesian computation was used for parameter inference of experimental data from simulated data. For this we employed the R package abc v2.1 (Csillery et al., 2010). In short, a set of simulation parameters, *θ_i_*, is sampled from the prior distribution. This set of parameters corresponds to a set of simulated summary statistics, *S(y_i_)*, in this case phylogenetic tree shapes, which can be compared to the set of experimental summary statistics, *S(y_o_).* The Euclidean distance between the experimental and simulated summary statistics can then be calculated (*d(S(y_i_),S(y_o_)).* A threshold, *T,* is then selected—0.05 in our case—which rejects the lower 1-*T* sets of simulation parameters that correspond. The remaining parameters represent those that gave summary statistics with the highest similarity to the experimental summary statistics. These represent the posterior distribution of accepted parameters.

### Cell cultivation procedures

Cal51 cells expressing stably integrated RFP-tagged histone H2B and GFP-tagged *α*-tubulin were generated as previously described (Zasadil et al., 2014). Cells were maintained at 37°C and 5% CO_2_ in a humidified, water-jacketed incubator and propagated in Dulbecco’s Modified Eagle’s Medium (DMEM) – High Glucose formulation (Cat #: 11965118) supplemented with 10% fetal bovine serum and 100 units/mL penicillin-streptomycin. Paclitaxel (Tocris Bioscience, Cat #: 1097/10) used for cell culture experiments was dissolved in DMSO.

### Time-lapse fluorescence microscopy

Cal51 cells were transduced with lentivirus expressing mNeonGreen-tubulin-P2A-H2B-FusionRed. A monoclonal line was treated with 20 nM paclitaxel for 24, 48, or 72 hours before timelapse analysis at 37°C and 10% CO_2_. Five 2 µm z-plane images were acquired using a Nikon Ti-E inverted microscope with a cMos camera at 3-minute intervals using a 40X/0.75 NA objective lens and Nikon Elements software.

### Flow cytometric analysis and cell sorting

Cells were harvested with trypsin, passed through a 35 μm mesh filter, and rinsed with PBS prior to fixation in ice cold 80% methanol. Fixed cells were stored at −80°C until analysis and sorting at which point fixed cells were resuspended in PBS containing 10 μg/ml DAPI for cell cycle analysis

*<u>Flow cytometric analysis</u>.* Initial DNA content and cell cycle analyses were performed on a 5 laser BD LSR II. Doublets were excluded from analysis via standard FSC/SSC gating procedures. DNA content was analyzed via DAPI excitation at 355 nm and 450/50 emission using a 410 nm long pass dichroic filter.

*<u>Fluorescence activated cell sorting</u>*. Cell sorting was performed using the same analysis procedures described above on a BD FACS AriaII cell sorter. In general, single cells were sorted through a 130-μm low-pressure deposition nozzle into each well of a 96-well PCR plate containing 10 μl Lysis and Fragmentation Buffer cooled to 4°C on a Eppendorf PCR plate cooler. Immediately after sorting PCR plates were centrifuged at 300 x g for 60 seconds. For comparison of single-cell sequencing to bulk sequencing, 1000 cells were sorted into each ‘bulk’ well. The index of sorted cells was retained allowing for the *post-hoc* estimation of DNA content for each cell.

### Low-coverage single-cell whole genome sequencing

Initial library preparation for low-coverage scDNAseq was performed as previously described (Leung et al., 2016) and adapted for low coverage whole genome sequencing instead of high coverage targeted sequencing. Initial genome amplification was performed using the GenomePlex® Single Cell Whole Genome Amplification Kit and protocol (Sigma Aldrich, Cat #: WGA4). Cells were sorted into 10 μl pre-prepared Lysis and Fragmentation buffer containing Proteinase K. DNA was fragmented to an average of 1 kb in length prior to amplification. Single cell libraries were purified on a 96-well column plate (Promega, Cat #: A2271). Library fragment distribution was assessed via agarose gel electrophoresis and concentrations were measured on a Nanodrop 2000. Sequencing libraries were prepared using the QuantaBio sparQ DNA Frag and Library Prep Kit. Amplified single-cell DNA was enzymatically fragmented to ∼250 bp, 5’-phosphorylated, and 3’-dA-tailed. Custom Illumina adapters with 96 unique 8 bp P7 index barcodes were ligated to individual libraries to enable multiplexed sequencing (Leung et al., 2016). Barcoded libraries were amplified following size selection via Axygen™AxyPrep Mag™ beads (Cat #: 14-223-152). Amplified library DNA concentration was quantified using the Quant-iT™ Broad-Range dsDNA Assay Kit (Thermo, Cat #: Q33130). Single-cell libraries were pooled to 15 nM and final concentration was measured via qPCR. Single-end 100bp sequencing was performed on an Illumina HiSeq2500.

### Single-cell copy number sequencing data processing

Single-cell DNA sequence reads were demultiplexed using unique barcode index sequences and trimmed to remove adapter sequences. Reads were aligned to GRCh38 using Bowtie2. Aligned BAM files were then processed using Ginkgo to make binned copy number calls. Reads are aligned within 500kb bins and estimated DNA content for each cell, obtained by flow cytometric analysis, was used to calculate bin copy numbers based on the relative ratio of reads per bin (Garvin et al., 2015). We modified and ran Ginkgo locally to allow for the analysis of highly variable karyotypes with low ploidy values (see Code and Data Availability). Whole-chromosome copy number calls were calculated as the modal binned copy number across an individual chromosome. Cells with fewer than 100,000 reads were filtered out to ensure accurate copy number calls (Baslan et al., 2015). Cells whose predicted ploidy deviated more than 32% from the observed ploidy by FACS were also filtered out. The final coverage for the filtered dataset was 0.03 (Supplemental Figure 9).

### Review and approximation of mis-segregation rates from published studies

We reviewed the literature to extract per chromosome rates of mis-segregation for cell lines and clinical samples. Some studies publish these rates. For those that did not, we estimated these rates by approximating the plotted incidence of segregation errors thusly:

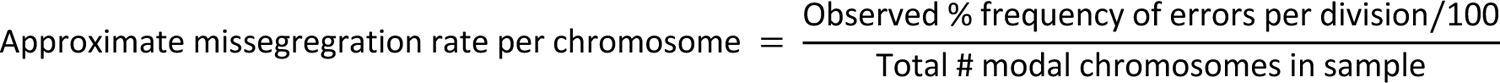

Modal chromosome numbers were either taken from ATCC where available or were assumed to equal 46. Observed % frequencies were approximated from published plots. Approximated rates assume that 1 chromosome is mis-segregated at a time.

## ABBREVIATIONS

ABC: approximate Bayesian computation

CIN: chromosomal instability

CFS: contextual chromosome fitness score

scDNAseq: single cell DNA sequencing

TOE scores: Tumor-Oncogene-Essential gene scores

WGD: whole genome doubling

## ACKNOWLEDGMENTS

This study was supported by grants to M.E.B. and B.A.W. from the NCI (5R01CA234904). A.R.L. was supported by the UW Cellular and Molecular Pathology (5T32GM081061) and the UW Genomic Sciences Training Program (5T32HG002760) NIH training grants. N.L.A. was supported by T32GM008692. A.S.Z. was supported in part by T32GM008688.Technical support comes from University of Wisconsin Carbone Cancer Center (UWCCC) Shared Resources funded by the UWCCC Support Grant P30 CA014520 – Flow Cytometry Core Facility (1S10RR025483-01), Cancer Informatics Shared Resource, Small Molecule Screening Facility. The authors thank the UW Biotechnology Center DNA Sequencing Facility for providing Illumina sequencing services. Special thanks go to Drs. Ana Bolhaqueiro, Bas Ponsioen, and Geert Kops for the provision of scDNAseq data for our analyses and to Dr. Caitlin Pepperell for valuable comments related to approximate Bayesian computation.

## AUTHOR CONTRIBUTIONS

Conception and design: A.R.L., B.A.W., M.E.B.

Development of methodology: A.R.L, N.L.A., M.E.B

Acquisition of data: A.R.L, A.S.Z

Analysis and interpretation of data: A.R.L, N.L.A., B.A.W, M.E.B

Writing, review, and revision of manuscript: A.R.L, A.S.Z, B.A.W, M.E.B

Administrative, technical, or material support: B.A.W, M.E.B

Study supervision: M.E.B

## COMPETING INTERESTS

The authors declare no competing interests.

## CODE AND DATA AVAILABILITY

DNA sequencing data from this study will be deposited in the NCBI Sequence Read Archive prior to final publication. Source code and data for modeling and analysis will be available on Open-Science Framework (OSF) prior to final publication.

## SUPPLEMENTAL FIGURE LEGENDS

Mean percent accuracy of ABC-inferred rates of mis-segregation due to paclitaxel taken from each set of 5 random samples using the observed rate of mis-segregation as the ‘true value’. Calculated as [Inline]. Dashed lines represent 80% and 90% accuracy. (D) Standard error of ABC-inferred rates of mis-segregation for each set of random samples from paclitaxel-treated cells. (E) ABC-inferred mis-segregation rates by sample size from simulations with known parameters (n = 5 samples). Points represent mean ± standard error across 5 samples for each of 9 selective pressure (S) values. Solid line represents a perfect correlation. Inner dashed line represent ± 10% margin. Outer dashed line represents ± 20% margin. Simulation parameters: Pmisseg ≤ 0.022; steps = 60; Hybrid model. (F) Mean percent accuracy of ABC-inferred rates of mis-segregation in simulations (parameters in E) taken at various sample sizes. Grey lines represent the mean percent accuracy of 5 random samples for each sample size for the same simulated population (n = 63 simulations). The dashed line represents 90% accuracy. Calculated as described above but taking the known simulation parameter as the ‘true’ value. (G) Standard error of ABC-inferred rates of mis-segregation in simulations (parameters in E) taken at various sample sizes. Grey lines represent the standard error of 5 random samples for each sample size for the same simulated population (n = 63 simulations). (H) ABC-inferred mis-segregation rates by sample size from simulations with known parameters (n = 5 samples). Points represent mean ± standard error across 5 samples for each of 9 selective pressure (S) values. Solid line represents a perfect correlation. Inner dashed line represent ± 10% margin. Outer dashed line represents ± 20% margin. Simulation parameters: Pmisseg = [0.001..0.5]; steps = 1; Hybrid model. (I) Mean percent accuracy of ABC-inferred rates of mis-segregation in simulations (parameters in H) taken at various sample sizes. Grey lines represent the mean percent accuracy of 5 random samples for each sample size for the same simulated population (n = 261 simulations). The dashed line represents 90% accuracy. (J) Standard error of ABC-inferred rates of mis-segregation in simulations (parameters in H) taken at various sample sizes. Grey lines represent the standard error of 5 random samples for each sample size for the same simulated population (n = 261 simulations). Note: Red lines in F, G, I, and J represent the median.

**Supplemental Figure 1.**
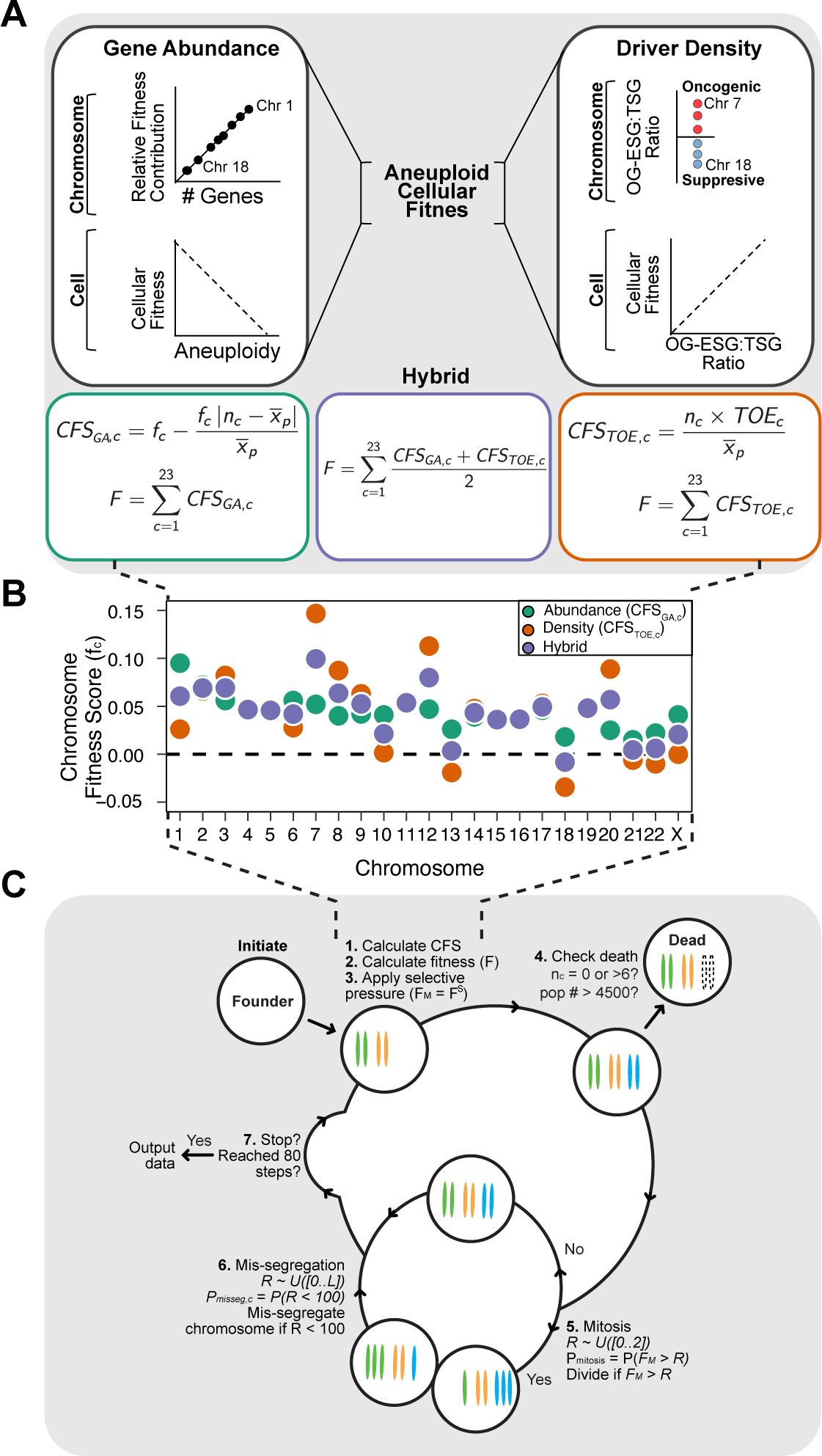
Expanded model of chromosome mis-segregation and karyotypic selection. (A) Models of selection on aneuploid karyotypes. **Left**. The Gene Abundance model dictates that chromosomes that encode a larger number of genes contribute more to cellular fitness (F). Thus, large chromosomes have a higher fitness score (fc). Deviation from the average ploidy of the population results in a reduced Contextual Fitness Score (CFS) for each chromosome, the sum of which represents the fitness of the cell. **Right**. The Driver Density Model dictates that the fitness contribution of a chromosome depends on the ratio of oncogenes and essential genes to tumor suppressors (OG-ESG:TSG). Gaining chromosomes with a higher OG-ESG:TSG ratio provides a fitness advantage while gaining more suppressive chromosomes invokes a fitness cost. These scores are still normalized to the ploidy of the average ploidy of the population to ensure that higher ploidy populations are not arbitrarily more fit. **Middle.** The Hybrid model takes the average of the fitness scores calculated in the other models. (B) Base chromosome fitness scores for each model. Only the Hybrid and Driver Density model have negatively scored chromosomes, meaning their loss provides a fitness benefit. (C) Populations are founded by 100 founder cells, unless otherwise stated, and the simulation is initiated. **1**. CFS values are calculated for each chromosome in a cell according to the chosen model. **2**. Cellular fitness is calculated based on CFS values. **3**. Selective pressure (S) is applied on cellular fitness values (F). **4**. Cells are checked to see if any death conditions are met and if the population limit is met. **5**. Cells probabilistically enter mitosis if their fitness value exceeds a random float (R) between 0 and 2. Thus Pmitosis = P(FM > R). If a cell does not enter mitosis, it skips the next step. **6**. If a cell enters mitosis, each chromosome has an opportunity to mis-segregate probabilistically. For each chromosome, a mis-segregation occurs if a random number (R), from 0 to some limit (L), falls below 100. Thus, Pmisseg,c = P(R<100). After chromosomes are mis-segregated, the cycle repeats and new CFS values are calculated, unless (**7**) stop conditions are met.

**Supplemental Figure 2.**
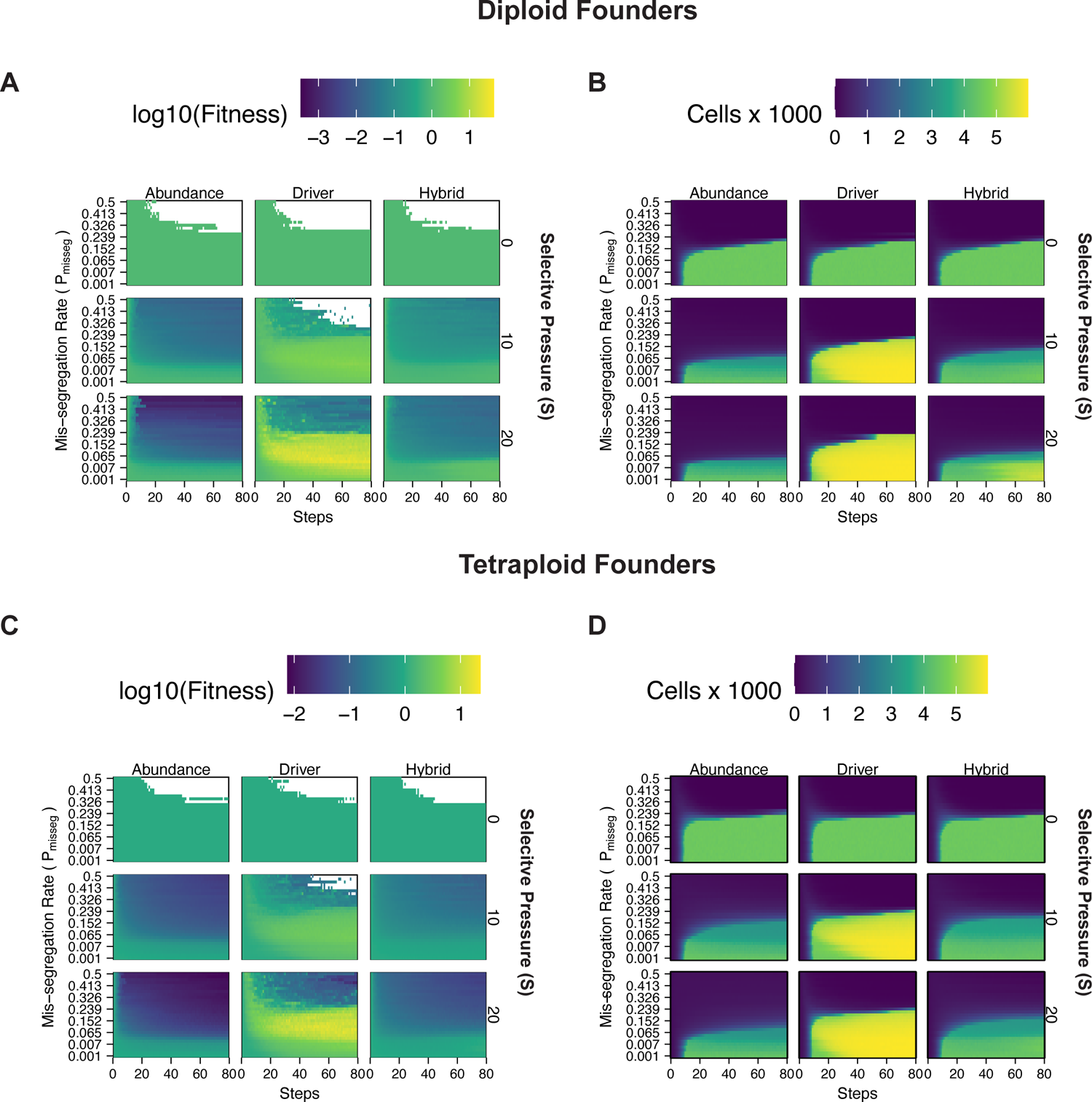
Fitness of diploid and tetraploid CIN+ populations (A) Fitness landscape of simulations founded by diploid cells. (B) Size of simulated populations founded by diploid cells. (C) Fitness landscape of simulations founded by tetraploid cells. (D) Size of simulated populations founded by tetraploid cells.

**Supplemental Figure 3.**
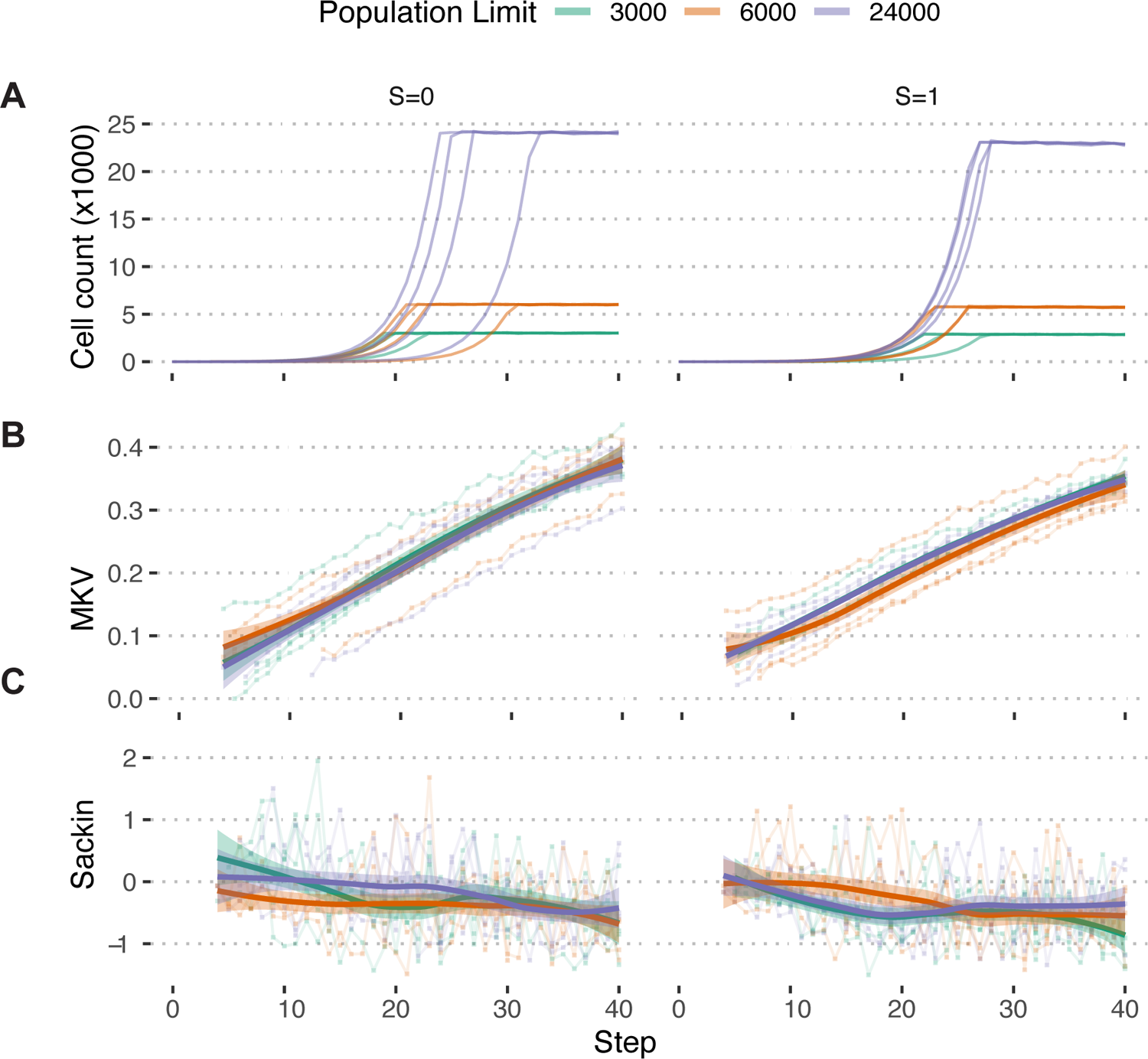
Population growth limits do not bias population measures (A) Growth curves of populations simulated under the Hybrid model with S = [0,1] and Pmisseg = 0.022 and limited to 3000, 6000, and 24000 cells (n = 4 simulations each). (B) MKV (normalized to mean ploidy of the population) values steadily increase over time. Loess regression curves show no significant deviations based on the population threshold, regardless of selection. (C) Sackin-Yule index values for each population over time. No significant deviations based on the population threshold, regardless of selection.

**Supplemental Figure 4.**
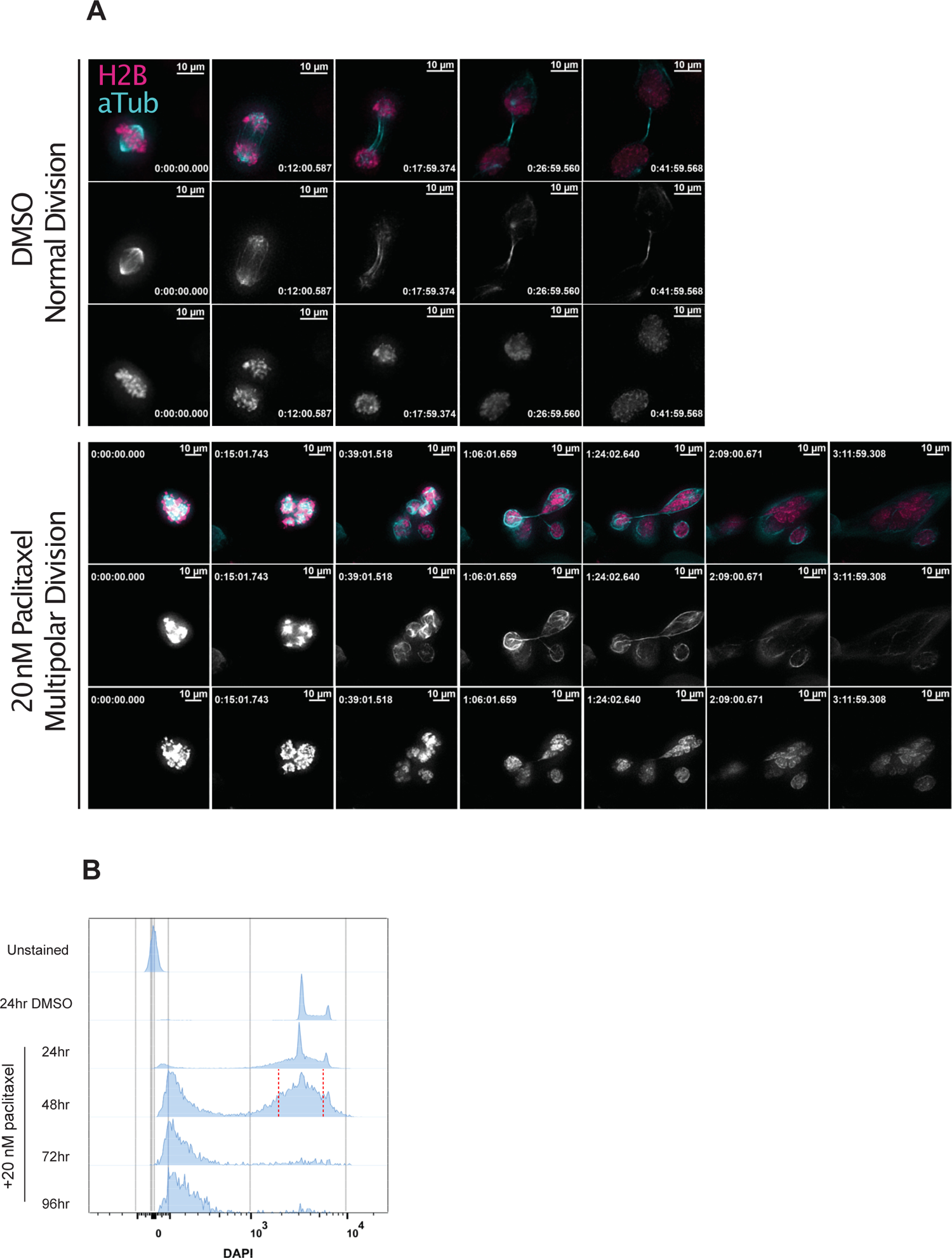
Induction of extensive chromosome mis-segregation via paclitaxel (A) Immunofluorescence time lapse montage of control Cal51 cells undergoing normal mitosis (top) and paclitaxel-treated treated cells undergoing a multipolar anaphase (middle) and partial cytokinesis failure (bottom). (B) Cell cycle profiles from flow cytometric analysis of Cal51 cells treated with either DMSO (72 hours) or 20 nM paclitaxel for 24, 48, or 72 hours. For FACS, cells treated for 48 hours were sorted into individual wells of 96 well plates. Sorting gate is shown by the red, dotted line.

**Supplemental Figure 5.**
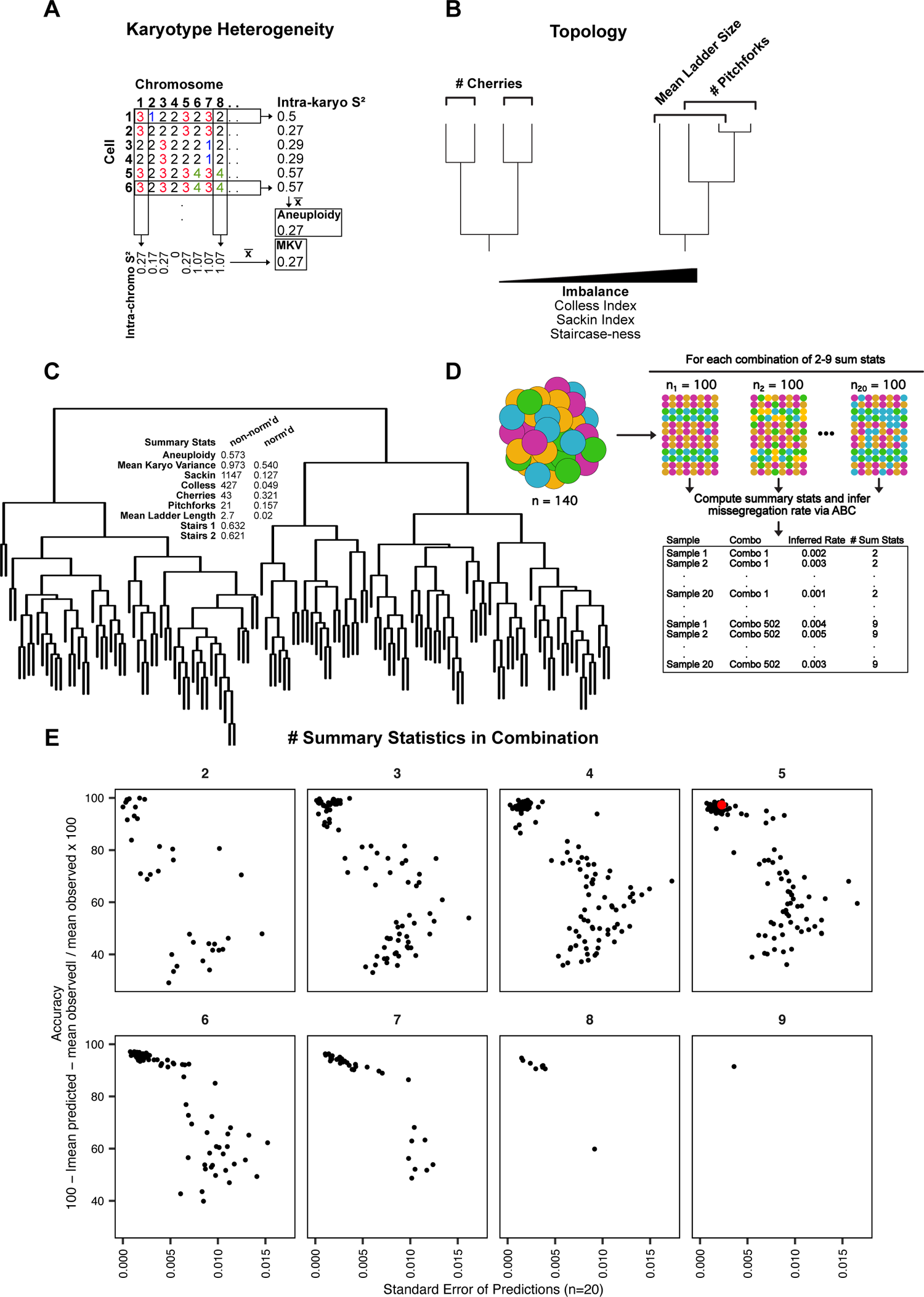
Summary statistic optimization for ABC (A) Example scheme for calculating aneuploidy and MKV. (B) Examples of phylogenetic topology metrics. (C) Phylogenetic reconstruction of a population of Cal51 cells treated with 20 nM paclitaxel for 48 hours and associated heterogeneity and topology metrics. Normalized and non-normalized summary statistics are displayed (see Materials & Methods). (D) Analytical scheme to identify most accurate and least variable combinations of heterogeneity and topology metrics. For each combination of 2-9 metrics, we iteratively re-sampled and remeasured the rate of missegregation in 100 random cells, 20 times, from our original dataset of paclitaxel-treated Cal51 cells. The red data point denotes our chosen combination for future analyses—average aneuploidy, MKV, Colless Index, Cherries, and Pitchforks. This combination contains both limits redundant measures (i.e. Colless and Sackin indices) and contains both heterogeneity and topology metrics. (E) Percent accuracy and standard error of the mean for 50 sampled measurements of 100 paclitaxel-treated cells from the original population, repeated for each combination of heterogeneity and topology measures.

**Supplemental Figure 6.**
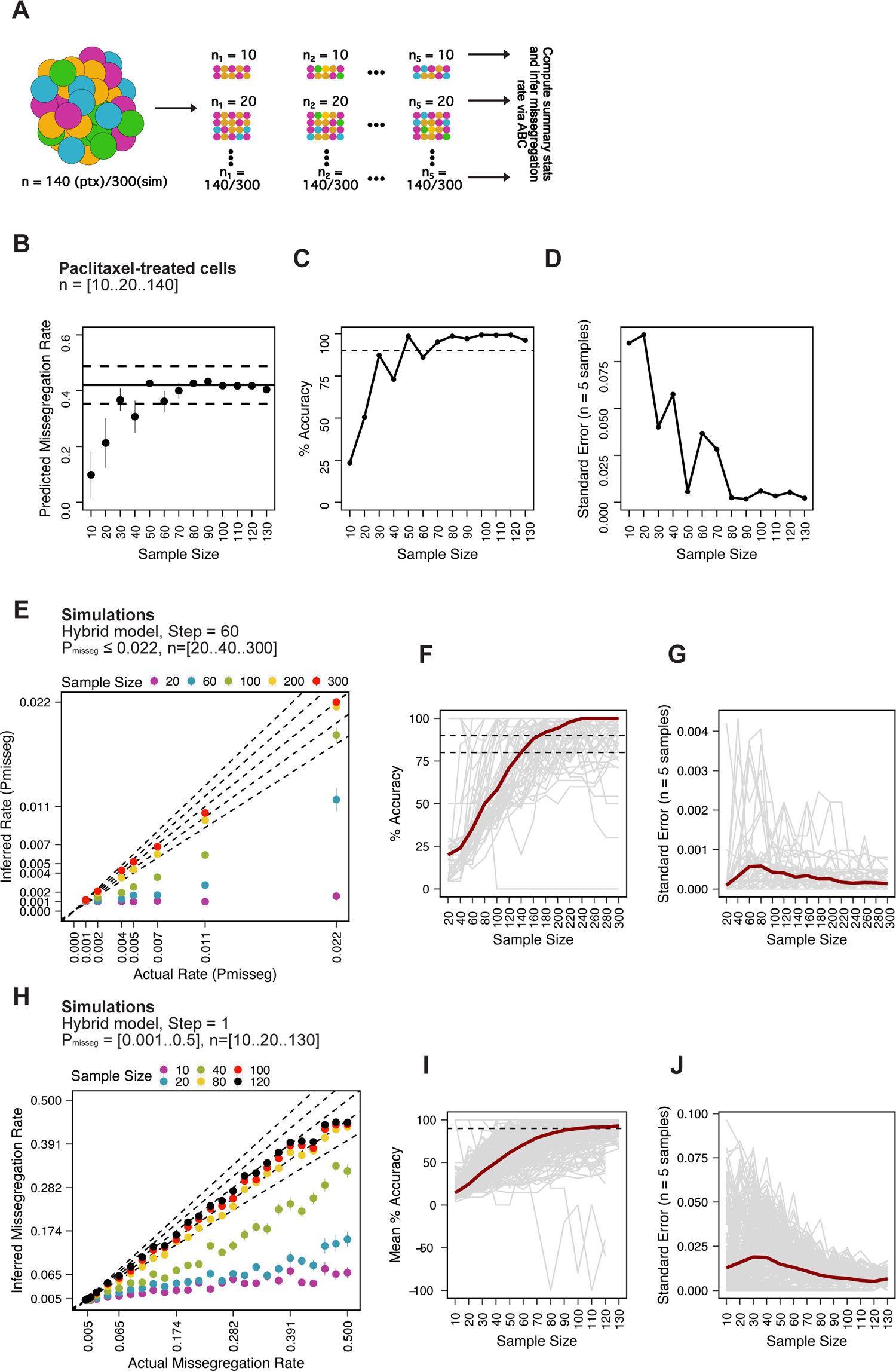
Minimum sampling of karyotype heterogeneity (A) Analytical scheme to optimize the number of cells to sample for measuring mis-segregation rates from karyotype heterogeneity. We iteratively re-sampled and remeasured the rate of mis-segregation for a range of sample sizes (n=5 random samples). (B) Predicted mis-segregation rates over a range of sample sizes (n=5 samples). Points and error bars are the mean ± standard error. Black solid line denotes the mean observed rate of mis-segregation induced by 20 nM paclitaxel. Black dashed lines are half the standard deviation of observed mis-segregation rates per cell. (C) true

**Supplemental Figure 7.**
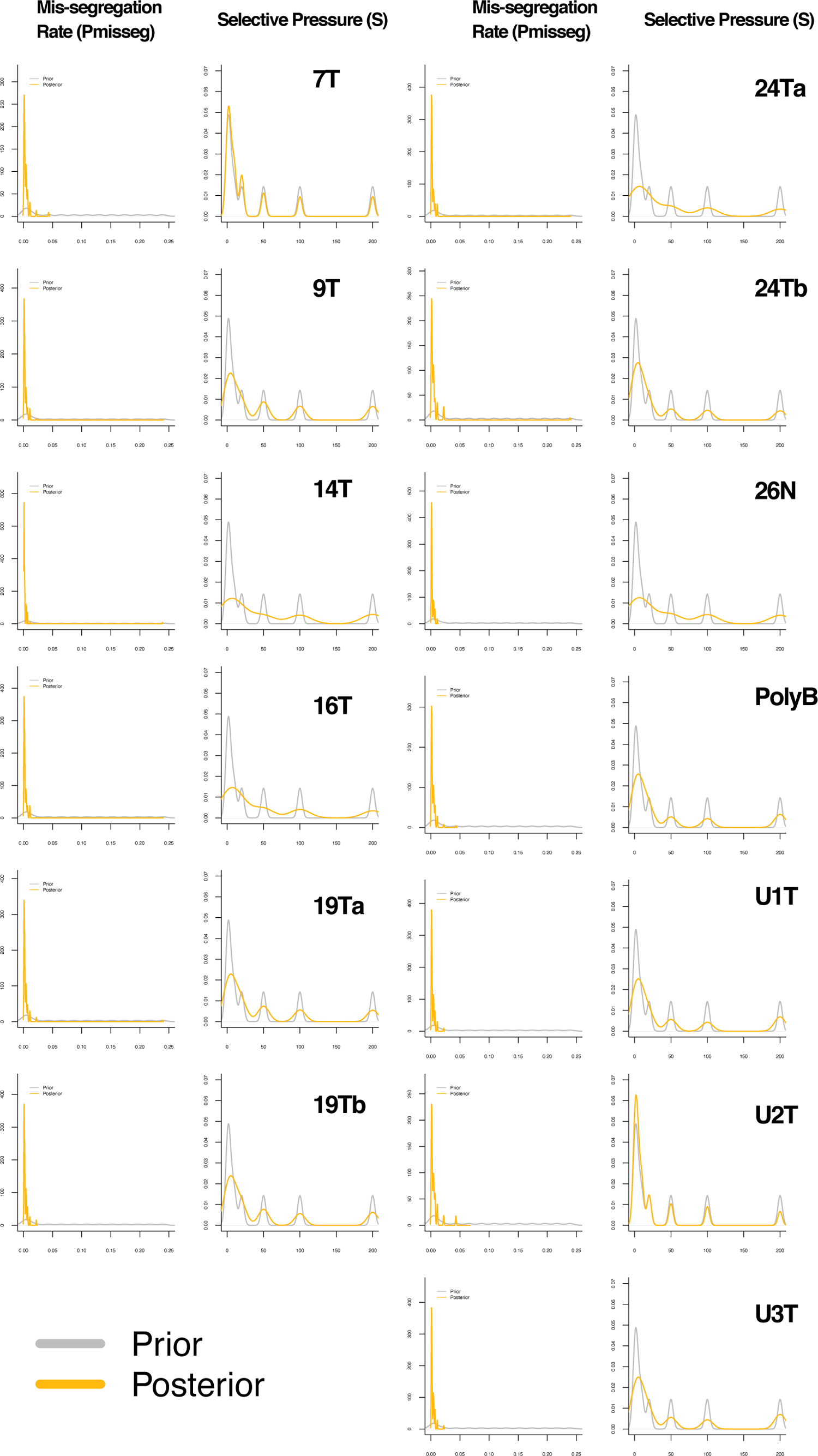
ABC-inferred mis-segregation rates and selective pressures in patient-derived samples Distributions of mis-segregation rates and selective pressures in patient-derived CRC organoids and a breast biopsy from Bolhaqueiro et al., 2019 and Navin et al., 2011 respectively. The prior (grey) distribution represents the parameters used for simulation while the posterior (yellow) distribution represents the parameters from simulations whose observed measurements were similar to the measurements taken from the patient-derived sample.

**Supplemental Figure 8.**
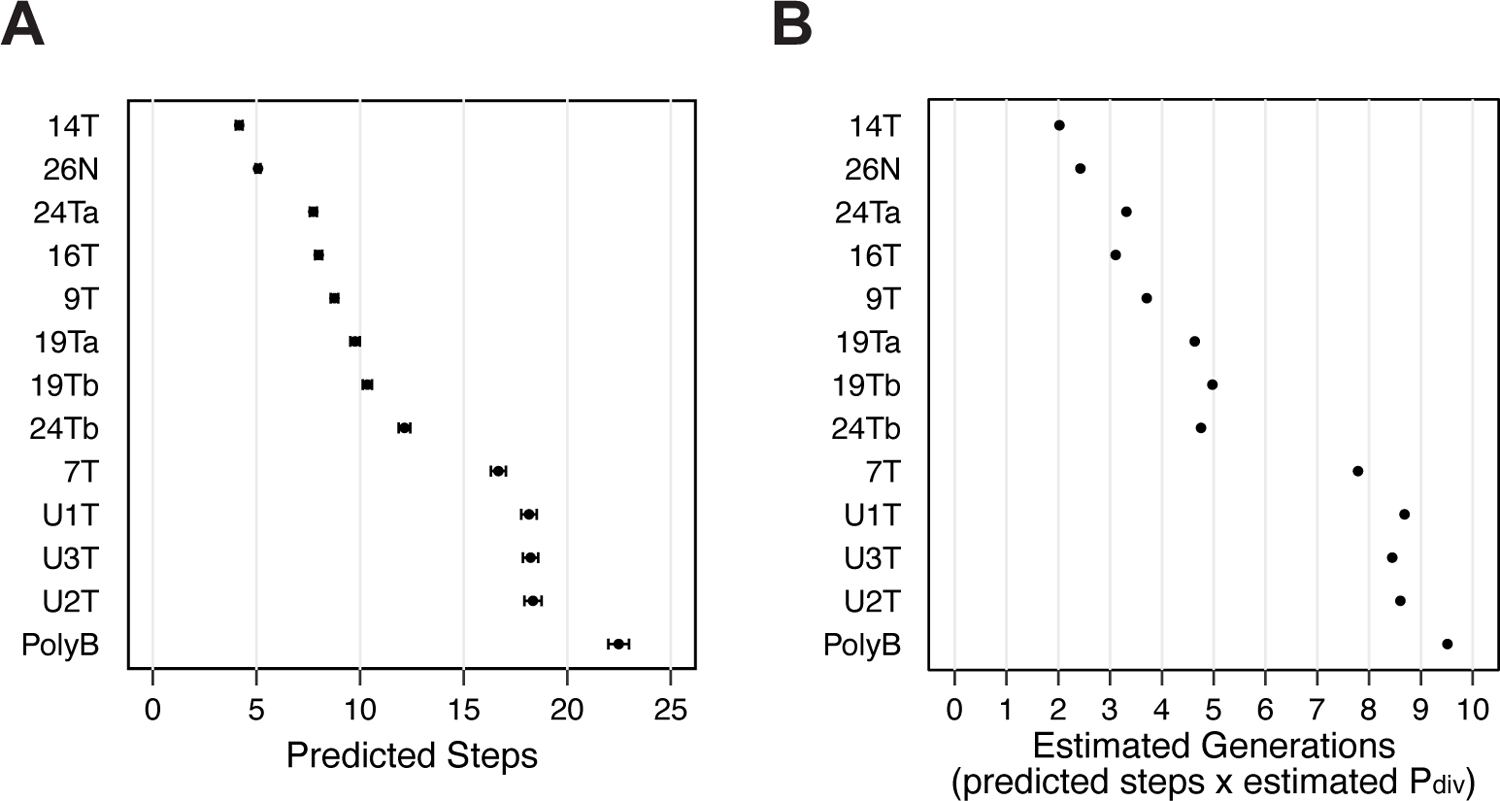
ABC-inferred step count in patient-derived samples (A) Number of steps experienced in each patient-derived sample, inferred via approximate Bayesian computation. (B) Inferred average number of generations experienced by cells in each patient-derived sample. The average Pdivision was calculated for each sample and then used to calculate the average number of generations using estimated generations = steps × P_div_

**Supplemental Figure 9.**
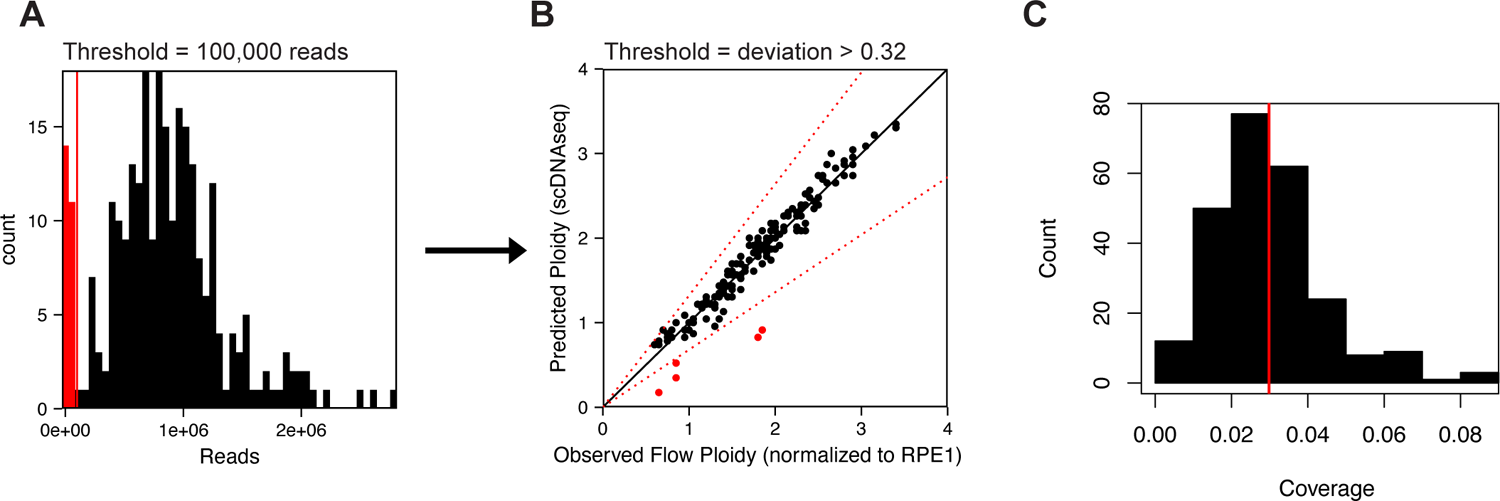
Single-cell sequencing QC (A) Read count for all sequencing samples and filtering threshold (red line and bins) of 100,000 reads. (B) Observed ploidy (FACS) vs predicted ploidy (scDNAseq). Predicted ploidy was calculated as the sum of copy number calls for each cell. Red dotted lines depict the 32% deviation threshold for filtering out poorly correlated cells. (C) Sequencing coverage for the final filtered dataset. The mean coverage (red line) was 0.03.

## SUPPLEMENTAL TABLES

**Supplemental Table 1.** Base chromosome-specific fitness scores for individual models

| CHR | Model |  |  |
| --- | --- | --- | --- |
|  | Gene Abundance | Driver Density | Hybrid |
| 1 | 0.095 | 0.026 | 0.0605 |
| 2 | 0.072 | 0.066 | 0.069 |
| 3 | 0.056 | 0.082 | 0.069 |
| 4 | 0.046 | 0.047 | 0.0465 |
| 5 | 0.048 | 0.043 | 0.0455 |
| 6 | 0.056 | 0.028 | 0.042 |
| 7 | 0.052 | 0.147 | 0.0995 |
| 8 | 0.040 | 0.087 | 0.0635 |
| 9 | 0.042 | 0.063 | 0.0525 |
| 10 | 0.041 | 0.002 | 0.0215 |
| 11 | 0.055 | 0.052 | 0.0535 |
| 12 | 0.047 | 0.113 | 0.08 |
| 13 | 0.026 | -0.019 | 0.0035 |
| 14 | 0.039 | 0.048 | 0.0435 |
| 15 | 0.034 | 0.038 | 0.036 |
| 16 | 0.036 | 0.037 | 0.0365 |
| 17 | 0.046 | 0.053 | 0.0495 |
| 18 | 0.018 | -0.034 | -0.008 |
| 19 | 0.047 | 0.049 | 0.048 |
| 20 | 0.025 | 0.089 | 0.057 |
| 21 | 0.015 | -0.006 | 0.0045 |
| 22 | 0.022 | -0.010 | 0.006 |
| X | 0.041 | 0 | 0.0205 |

**Supplemental Table 2.** Approximate reported per chromosome mis-segregation rates

| 1st Author | DOI | Model | Tumor? | Statistic | Assessment | Approximate observed frequency % | Approx modal chromosome # (ATCC) | Approximate mis-segregation rate (per chromosome) |
| --- | --- | --- | --- | --- | --- | --- | --- | --- |
| Bakhoun | https://doi.org/10.1158/1078-0432.CCR-11-2049 | Tumor-TMA | Tumor | Reported | Lagging/Bridging | 31.3 | 46 | 0.00680 |
| Orr | https://doi.org/10.1016/j.celrep.2016.10.030 | U2OS | Tumor | Approx. Mean | Lagging | 32.5 | 46 | 0.00707 |
| Orr | https://doi.org/10.1016/j.celrep.2016.10.030 | HeLa | Tumor | Approx. Mean | Lagging | 22 | 82 | 0.00268 |
| Orr | https://doi.org/10.1016/j.celrep.2016.10.030 | SW-620 | Tumor | Approx. Mean | Lagging | 22.5 | 50 | 0.00450 |
| Orr | https://doi.org/10.1016/j.celrep.2016.10.030 | RPE1 | Non-tumor | Approx. Mean | Lagging | 2.5 | 46 | 0.00054 |
| Orr | https://doi.org/10.1016/j.celrep.2016.10.030 | BJ | Non-tumor | Approx. Mean | Lagging | 8 | 46 | 0.00174 |
| Nicholson | https://doi.org/10.7554/eLife.05068 | Amniocyte | Non-tumor | Approx. Mean | Lagging | 0 | 46 | 0.00000 |
| Nicholson | https://doi.org/10.7554/eLife.05068 | DLD1 | Tumor | Approx. Mean | Lagging | 1 | 46 | 0.00022 |
| Dewhurst | https://doi.org/10.1158/2159-8290.CD-13-0285 | HCT116-Diploid | Tumor | Approx. Mean | Lagging/Bridging | 23 | 45 | 0.00511 |
| Dewhurst | https://doi.org/10.1158/2159-8290.CD-13-0285 | HCT116-Tetraploid | Tumor | Approx. Mean | Lagging/Bridging | 50 | 90 | 0.00556 |
| Bakhoun | https://doi.org/10.1038/ncb1809 | U2OS | Tumor | Reported | Lagging |  | 46 | 0.01000 |
| Zasadil | https://doi.org/10.1126/scitranslmed.3007965 | CAL51 | Tumor | Approx. Mean | Lagging | 0.5 | 44 | 0.00011 |
| Thompson | https://doi.org/10.1083/jcb.200712029 | RPE1 | Non-tumor | Approx. Mean | Acute aneuploidy via FISH |  | 46 | 0.00025 |
| Thompson | https://doi.org/10.1083/jcb.200712029 | HCT116-Diploid | Tumor | Approx. Mean | Acute aneuploidy via FISH |  | 45 | 0.00025 |
| Thompson | https://doi.org/10.1083/jcb.200712029 | HT29 | Tumor | Approx. Mean | Acute aneuploidy via FISH |  | 71 | 0.00250 |
| Thompson | https://doi.org/10.1083/jcb.200712029 | Caco2 | Tumor | Approx. Mean | Acute aneuploidy via FISH |  | 96 | 0.00900 |
| Thompson | https://doi.org/10.1083/jcb.200712029 | MCF-7 | Tumor | Approx. Mean | Acute aneuploidy via FISH |  | 82 | 0.00700 |
| Bakhoun | https://doi.org/10.1016/j.cub.2014.01.019 | HCT116-Diploid | Tumor | Approx. Mean | Lagging | 6 | 45 | 0.00133 |
| Bakhoun | https://doi.org/10.1016/j.cub.2014.01.019 | DLD1 | Tumor | Approx. Mean | Lagging | 2 | 46 | 0.00043 |
| Bakhoun | https://doi.org/10.1016/j.cub.2014.01.019 | HT29 | Tumor | Approx. Mean | Lagging | 14 | 71 | 0.00197 |
| Bakhoun | https://doi.org/10.1016/j.cub.2014.01.019 | SW-620 | Tumor | Approx. Mean | Lagging | 12 | 50 | 0.00240 |
| Bakhoun | https://doi.org/10.1016/j.cub.2014.01.019 | MCF-7 | Tumor | Approx. Mean | Lagging | 17 | 82 | 0.00207 |
| Bakhoun | https://doi.org/10.1016/j.cub.2014.01.019 | HeLa | Tumor | Approx. Mean | Lagging | 13 | 82 | 0.00159 |
| Worrall | https://doi.org/10.1016/j.celrep.2018.05.047 | BJ | Non-tumor | Approx. Mean | Unspecified Error | 5 | 46 | 0.00109 |
| Worrall | https://doi.org/10.1016/j.celrep.2018.05.047 | RPE1 | Non-tumor | Approx. Mean | Unspecified Error | 5 | 46 | 0.00109 |

## Notes

**Financial Support:** This work was supported by NIH grant 1R01CA234904 (M.E.B., B.A.W.) and P30CA014520. A.R.L was supported in part by 5T32GM081061-09 and 2T32HG002760-16. N.L.A. was supported by T32GM008692. A.S.Z. was supported in part by T32GM008688.

### Competing Interest Statement

M.E.B declares they are on the Medical Advisory Board of Strata Oncology, and that they receive research funding from Strata, Abbvie, Genetech, Puma, Arcus, and Loxo/Lilly. All other authors declare no financial conflicts of interest.

